# Associations between epilepsy-related polygenic risk and brain morphology in childhood

**DOI:** 10.1101/2025.01.17.633277

**Authors:** Alexander Ngo, Lang Liu, Sara Larivière, Valeria Kebets, Serena Fett, Clara F. Weber, Jessica Royer, Eric Yu, Raúl Rodríguez-Cruces, Zhiqiang Zhang, Leon Qi Rong Ooi, B. T. Thomas Yeo, Birgit Frauscher, Casey Paquola, Maria Eugenia Caligiuri, Antonio Gambardella, Luis Concha, Simon S. Keller, Fernando Cendes, Clarissa L. Yasuda, Leonardo Bonilha, Ezequiel Gleichgerrcht, Niels K. Focke, Raviteja Kotikalapudi, Terence J. O’Brien, Benjamin Sinclair, Lucy Vivash, Patricia M. Desmond, Elaine Lui, Anna Elisabetta Vaudano, Stefano Meletti, Reetta Kälviäinen, Hamid Soltanian-Zadeh, Gavin P. Winston, Vijay K. Tiwari, Barbara A. K. Kreilkamp, Matteo Lenge, Renzo Guerrini, Khalid Hamandi, Theodor Rüber, Tobias Bauer, Orrin Devinsky, Pasquale Striano, Erik Kaestner, Sean N. Hatton, Lorenzo Caciagli, Matthias Kirschner, John S. Duncan, Paul M. Thompson, ENIGMA Consortium Epilepsy Working Group, Carrie R. McDonald, Sanjay M. Sisodiya, Neda Bernasconi, Andrea Bernasconi, Ziv Gan-Or, Boris C. Bernhardt

## Abstract

Temporal lobe epilepsy with hippocampal sclerosis (TLE-HS) is associated with a complex genetic architecture, but the translation from genetic risk factors to brain vulnerability remains unclear. Here, we examined associations between epilepsy-related polygenic risk scores for HS (PRS-HS) and brain structure in a large sample of neurotypical children, and correlated these signatures with case-control findings in in multicentric cohorts of patients with TLE-HS. Imaging-genetic analyses revealed PRS-related cortical thinning in temporo-parietal and fronto-central regions, strongly anchored to distinct functional and structural network epicentres. Compared to disease-related effects derived from epilepsy case-control cohorts, structural correlates of PRS-HS mirrored atrophy and epicentre patterns in patients with TLE-HS. By identifying a potential pathway between genetic vulnerability and disease mechanisms, our findings provide new insights into the genetic underpinnings of structural alterations in TLE-HS and highlight potential imaging-genetic biomarkers for early risk stratification and personalized interventions.

## Introduction

Epilepsy is characterized by an enduring predisposition to recurrent spontaneous seizures and affects over 50 million people worldwide.^1^ One of the most common forms of epilepsy is temporal lobe epilepsy (TLE), a focal epilepsy associated pathologically with hippocampal sclerosis (HS) and pharmaco-resistance. Cumulative evidence has underscored the complexity of TLE-HS, revealing contributions of genetic and acquired factors in epileptogenesis. With seizure onsets typically in childhood and adolescence,^2^ developmental transitions spanning youth represent a key window for epilepsy risk. Adequately capturing the condition’s effects on brain organization, particularly in development, may advance our understanding of brain mechanisms giving rise to seizures and may have important implications for disease monitoring and early diagnosis.

In addition to its typical association with mesiotemporal pathology, neuroimaging evidence in patients with TLE-HS has identified widespread structural alterations. Magnetic resonance imaging (MRI) analysis of brain morphology has established robust structural compromise in the hippocampus, subcortical regions as well as more widespread temporal and fronto-central cortical systems. These findings were initially shown in single centre studies,^3–6^ and more recently confirmed in large-scale multisite initiatives, notably ENIGMA-Epilepsy.^7,8^ The latter initiative has mapped consistent patterns of multilobar atrophy in TLE-HS, and furthermore contextualized findings with measures of brain network architecture confirming temporo-limbic regions as epicentres of distributed structural pathology.^9^ Despite a likely influence of environmental factors and clinical events on brain structure in TLE,^10^ there has been growing evidence of important genetic influence,^11^ suggesting a possible mechanism contributing to this classical disease phenotype.

Epilepsy has a complex genetic architecture, with many contributory genetic factors.^12–16^ Variants underlying many different monogenic forms of epilepsy are rare, yet of large effect that can confer high risk or be causally responsible for the disease.^17,18^ Despite the clinical implications of these variants, common epilepsy syndromes, particularly TLE-HS, rarely carry such variants and presumably have a complex, multigenic inheritance.^19^ Causation may therefore be attributable to the synergy of multiple genetic variants interacting with each other, together with acquired environmental factors. Recent genome-wide association studies (GWAS) have identified common risk alleles.^13–16^ These individual genetic risk variants are usually of small effect and cannot quantify risk or inform prognosis and treatment.^20^ However, genome-wide profiling using polygenic risk scores (PRS) may provide a window into the genetic liability of the disease. By estimating the combined effect of individual single nucleotide polymorphisms (SNPs), it can collectively capturing the variance explained by these common alleles and provide an individualized measure of genetic risk.^21–23^ While previous studies have revealed enriched genetic vulnerability for epilepsy in patients,^24–26^ the consequences of epilepsy susceptibility on disease phenotypes, such as brain morphology, have not been systematically charted. Investigating this micro-to-macroscale mechanism may provide insight into the translation of genetic vulnerability to disease etiology or consequences.

In this study, we aimed to uncover the cumulative effects of epilepsy-related genetic risk variants on structural brain organization during development. We analyzed structural MRI and genotyping data in a large population-based cohort of neurotypical children from the Adolescent Brain Cognitive Development study (ABCD).^27^ To investigate associations between genetic risk factors for epilepsy-related HS and brain-wide morphology, we generated PRS-related models of cortical thickness and subcortical volume. Network contextualization further identified connectome epicentres of PRS-HS effects—network pathways that may govern the genetically affected morphological patterning. To pinpoint common processes between genetic risk and disease pathologies, we employed spatial correlations with autocorrelation preserving null models and related structural effects of PRS-HS to disease-related atrophy and epicentres derived from large multi-site MRI-based datasets of patients and controls.^28–30^

## Results

### Genetic and neuroimaging data samples

We studied genetic and imaging data of 3,826 unrelated neurotypical children (mean ± standard deviation [SD] age = 10.0 ± 0.6 years; 2,052 males) from the multi-site ABCD 2.0.1 release.^31^ Epilepsy-related PRS was calculated to determine an individual’s genetic burden for TLE-HS. Based on previous GWAS summary statistics,^15^ we took the weighted sum of disease-related risk alleles in an individual’s genome, with weights reflecting the effect sizes of each variant. In parallel, cortical thickness across 68 gray matter brain regions and volumetric data from 12 subcortical gray matter regions and bilateral hippocampi were obtained from all children.^32^

### Structural correlates of PRS-HS

Imaging-genetic correlations based on surface-based linear models related PRS-HS to brain structure. We observed a significant and negative association between global cortical thickness and with genetic vulnerability (left hemisphere: *r* = -0.041, *p*_FDR_ < 0.05; right hemisphere: *r* = - 0.044, *p*_FDR_ < 0.05; **Figure 1A**). Adopting a regional approach, these effects colocalized to bilateral temporal pole and postcentral gyrus, left precuneus, inferior parietal and lateral occipital regions as well as right superior, and middle temporal, precentral and paracentral gyri (range *r* = -0.0501 – -0.0362, p_FDR_ < 0.05; **Figure 1B**).

**Figure 1.**
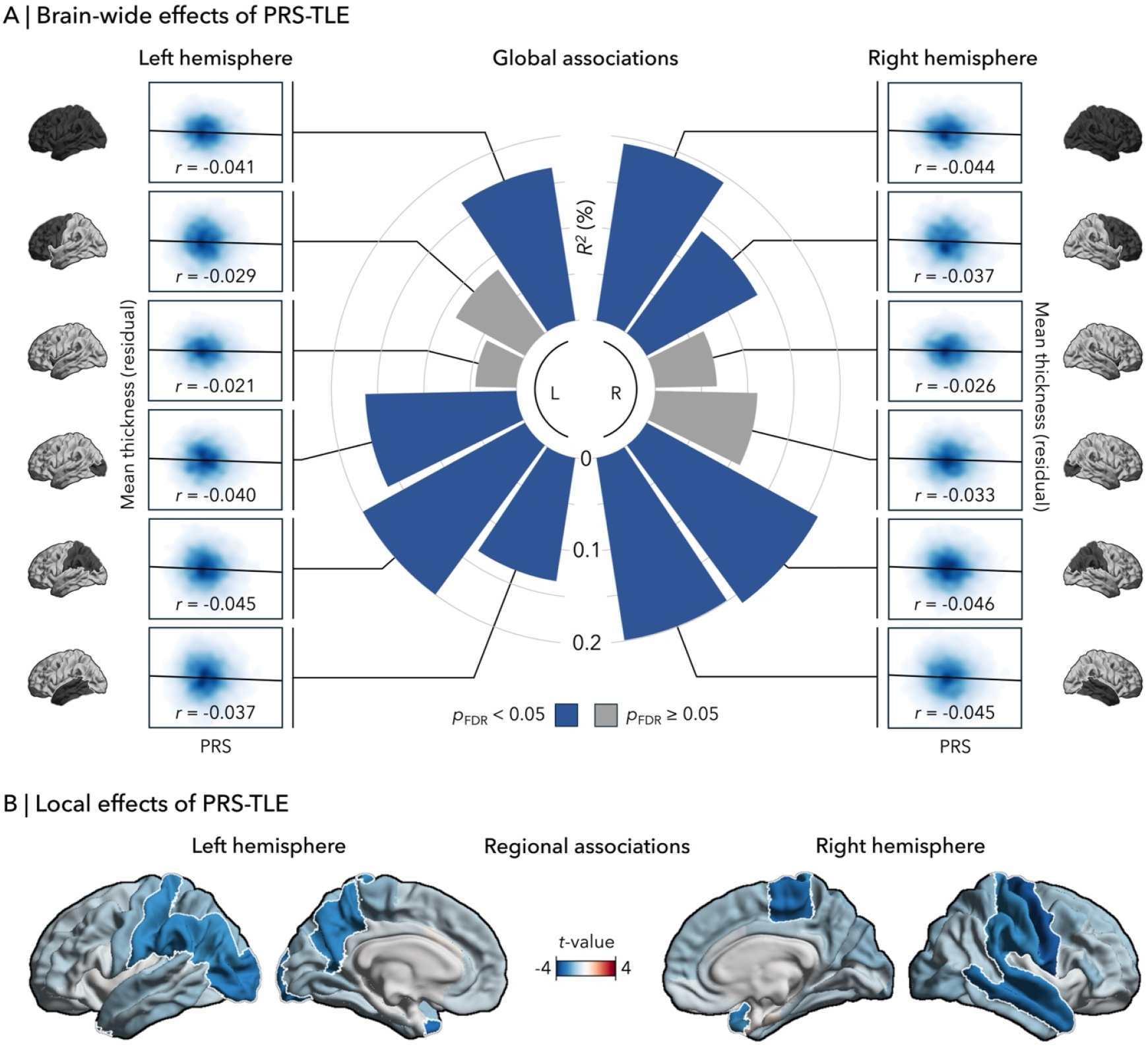
PRS-HS associations with cortical thickness (ABCD). **(A)** Distribution of genetic risk effects on morphology across the different lobes (in order from top to bottom: all, frontal, limbic, occipital, parietal, temporal). **(B)** Regional imaging-genetic correlations between PRS-HS and thickness. Blue and red colours represent negative and positive correlations, respectively. White outline indicates *p*_FDR_ < 0.05. L, left; PRS-HS; polygenic risk score for epilepsy-related hippocampal sclerosis; R, right.

After correcting for multiple comparisons, there were no significant relationships between PRS-HS and subcortical and hippocampal volume (all *p*_FDR_ ≥ 0.05; **Supplementary figure 1**).

### Network substrates of PRS-related structural changes

Given the large-scale effects of PRS-HS on cortical thickness, contextualizing imaging-genetic correlations with connectome architecture may provide insight into how localized genetic susceptibility propagates through distributed brain networks and predicts structural vulnerabilities. Data-driven epicentre mapping can identify one or more specific regions—or epicentres—whose connectivity profile spatially resembles structural effects of PRS-HS.^9,33–35^

We systematically correlated the functional and structural connections of each cortical and subcortical region to the imaging-genetic patterns (see *Figure 1*), with non-parametric spin permutation null models to control for spatial autocorrelation (5,000 repetitions, *p*-values were denoted as *p*_spin_).^36^ This analysis implicated bilateral temporal-limbic and parietal cortices, amygdalae, hippocampi, and thalami as the most significant functional and structural epicentres (all *p*_spin_ < 0.05; **Figure 2B**).

**Figure 2.**
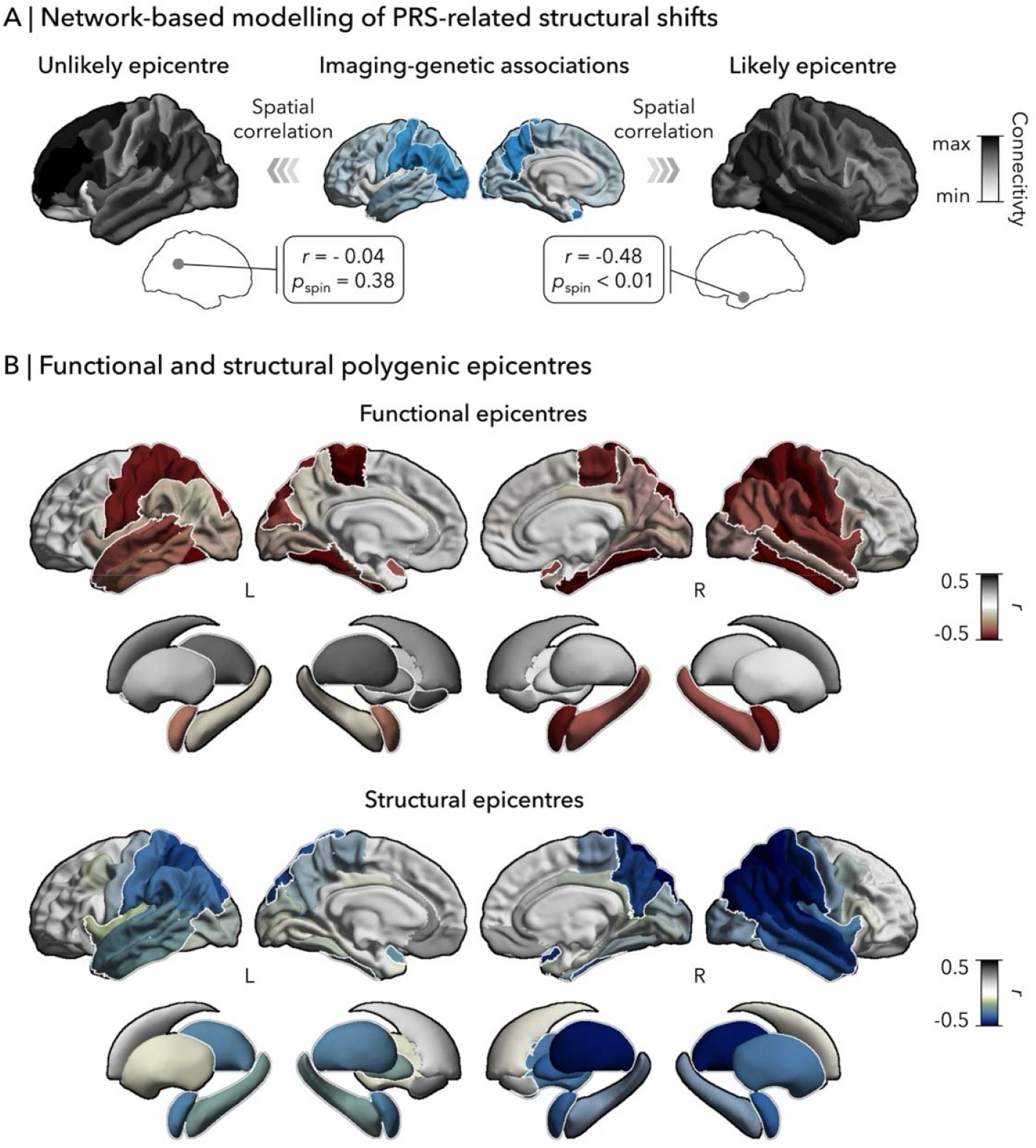
Network epicentres of morphological changes associated with PRS-HS. **(A)** Schematic representation of epicentre mapping approach using seed-based cortico- and subcortico-cortical connectivity. PRS-HS, polygenic risk score for epilepsy-related hippocampal sclerosis. **(B)** Correlation coefficients indexing spatial similarity between imaging-genetic effects and seed-based functional (*top*) and structural (*bottom*) connections for every cortical and subcortical region. Red and blue colours represent negative associations, while grey depicts positive correlations. White outline indicates *p*_spin_ < 0.05. L, left; R, right.

### Relation to epilepsy-specific atrophy and network epicentres

To link genetic vulnerability to disease alterations, we examined the spatial resemblance between imaging-genetic findings to atrophy patterns observed in patients with TLE-HS. We leveraged the ENIGMA-Epilepsy Consortium aggregating case-control MRI data of 785 individuals with TLE-HS and 1,512 healthy controls from multiple sites around the world.^8,28^ Between-group differences (ENIGMA-Epilepsy) revealed profound atrophy in patients, with strongest effects in bilateral precuneus, precentral, paracentral, and temporal cortices (*p*_FDR_ < 0.05; **Figure 3A**). Correlating alteration maps with PRS effects (from ABCD, see *Figure 1*) showed significant overlap with left (*r* = 0.63, *p*_spin_ = 0.001) and right TLE-HS (*r* = 0.59, *p*_spin_ = 0.0006; **Figure 3B**).

**Figure 3.**
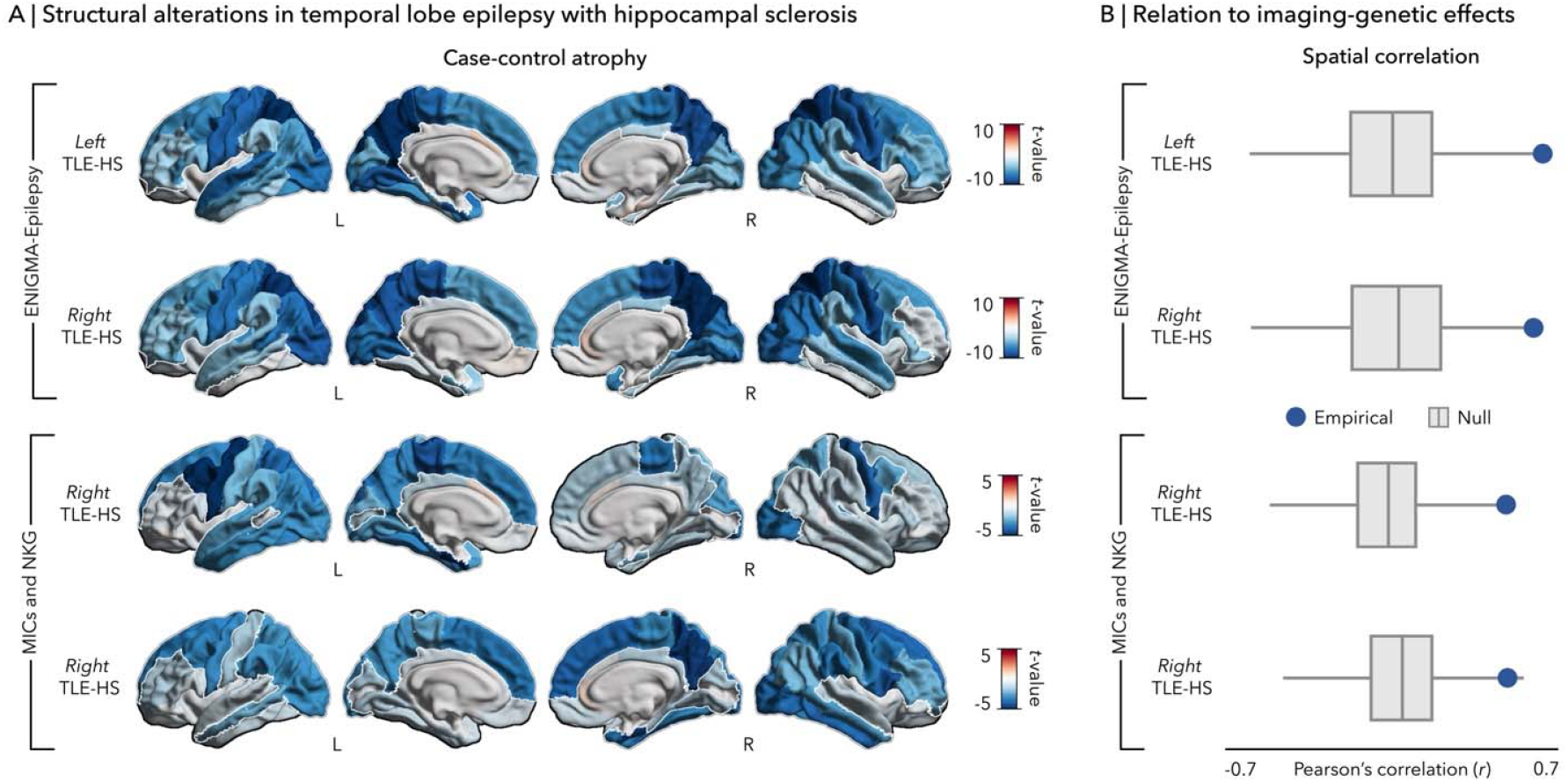
Comparison between PRS-HS effects and epilepsy case-control atrophy. **(A)** Case-control differences in left and right TLE-HS from ENIGMA-Epilepsy (*top*) and from MICs and NKG (*bottom*). Blue and red colours point to atrophy and hypertrophy in patients relative to healthy controls, respectively. Outline in white represents *p*_FDR_ < 0.05. L, left; R, right; TLE-HS, temporal lobe epilepsy with hippocampal sclerosis. **(B)** Spatial correlations between epilepsy-related atrophy (*top*: ENIGMA-Epilepsy; *bottom*: MICs and NKG) and imaging-genetic effect maps (ABCD) are compared against permutation-based null correlations. Points represent the empirical correlation (with significance defined as *p*_spin_ < 0.05). In the boxplots, the ends of boxes represent the first (25%) and third (75%) quartiles, the centre line (median) represents the second quartile of the null distribution (*n* = 5,000 permutations), the whiskers represent the non-outlier endpoints of the distribution.

Network mapping of TLE-related atrophy (ENIGMA-Epilepsy) revealed significant temporo-limbic and parieto-occipital epicentres in TLE-HS (*p*_FDR_ < 0.05; **Figure 4A**). Similarly, imaging-genetic epicentres (from ABCD, see *Figure 2*) were strongly correlated with disease epicentres in left TLE-HS (functional: *r* = 0.95, *p*_spin_ < 0.001; structural: *r* = 0.78, *p*_spin_ < 0.001), right TLE-HS (functional: *r* = 0.93, *p*_spin_ < 0.001; structural: *r* = 0.94, *p*_spin_ < 0.001; **Figure 4B**), suggesting potential pathway convergence between PRS-HS and TLE-HS effects.

**Figure 4.**
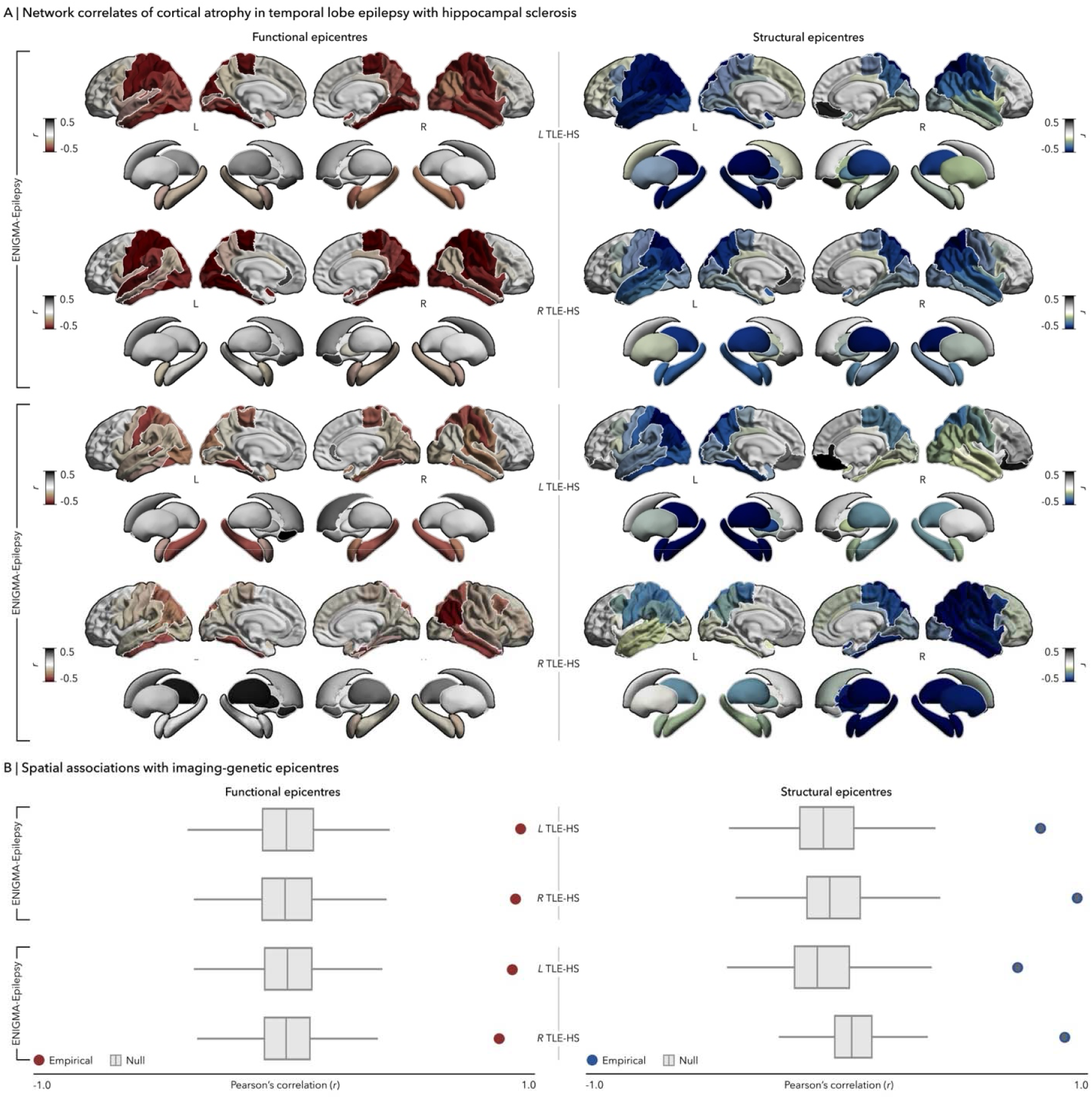
Comparison between imaging-genetic and epilepsy-related disease epicentres. **(A)** Functional and structural disease epicentres in left and right TLE-HS from ENIGMA-Epilepsy (*top*) and from MICs and NKG (*bottom*). Red and blue colours represent negative associations, while grey depicts positive correlations. Outline in white represents *p*_spin_ < 0.05. L, left; R, right; TLE-HS, temporal lobe epilepsy with hippocampal sclerosis. **(B)** Spatial correlations between epilepsy-related epicentres (*top*: ENIGMA-Epilepsy; *bottom*: MICs and NKG) and imaging-genetic effect maps (ABCD) are compared against permutation-based null correlations. Points represent the empirical correlation (with significance defined as *p*_spin_ < 0.05). In the boxplots, the ends of boxes represent the first (25%) and third (75%) quartiles, the centre line (median) represents the second quartile of the null distribution (*n* = 5,000 permutations), the whiskers represent the non-outlier endpoints of the distribution.

Assessing the consistency of these correlations, we repeated correlation analyses with a separate independent case-control set, collected from (*a*) Montreal Neurological Institute and Hospital (MICs; *n*_TLE-HS/HC_ = 23/36)^29^ and (*b*) Jinling Hospital (NKG; *n*_TLE-HS/HC_ = 37/57).^30^ we compared between PRS effects (from ABCD, see *Figure 1*) and disease-related atrophy (from MICs and NKG; **Figure 3A**) and observed moderate and highly significant positive correlations for left (*r* = 0.50, *p*_spin_ = 0.0002) and right TLE-HS (*r* = 0.41, *p*_spin_ = 0.009). Imaging-genetic epicentres (from ABCD, see *Figure 2*) were strongly similar with disease epicentres (**Figure 3A**) in left (functional: *r* = 0.93, *p*_FDR_ < 0.001; structural: *r* = 0.77, *p*_FDR_ < 0.001) and right TLE-HS (functional: *r* = 0.89, *p*_FDR_ < 0.001; structural: *r* = 0.89, *p*_FDR_ < 0.001; **Figure 3B**).

To further evaluate specificity of PRS-HS effects, we cross-referenced our imaging-genetic patterns (from ABCD, see *Figure 1* and *2*) with atrophy and disease epicentre maps in idiopathic generalized epilepsy (IGE), another common epilepsy syndrome, and six psychiatric disorders (attention deficit disorder [ADHD], autism spectrum disorder [ASD], bipolar disorder [BD], major depressive disorder [MDD], obsessive-compulsive disorder [OCD], and schizophrenia [SCZ]), all acquired from the ENIGMA Consortium.^28,37^ Spatial correlations between PRS-HS and TLE-HS effects (see *Figure 3* and *4*) were among the highest even when compared against the different conditions (**Table 1**; IGE: **Supplementary figure 2**; psychiatric conditions: **Supplementary figure 3** and **4**).

**Table 1.** Spatial correlation between effects of PRS-HS and different conditions.

| Condition | Analysis | Correlation ($r$) | P-value ($p_{\text{spin}}$) |
| --- | --- | --- | --- |
| IGE | Regional | 0.42 | 0.004 |
| | Functional epicentre | 0.76 | $< 0.001$ |
| | Structural epicentre | 0.70 | $< 0.001$ |
| ADHD | Regional | -0.29 | 0.071 |
| | Functional epicentre | -0.80 | $< 0.001$ |
| | Structural epicentre | -0.67 | $< 0.001$ |
| ASD | Regional | 0.14 | 0.322 |
| | Functional epicentre | 0.84 | $< 0.001$ |
| | Structural epicentre | 0.71 | $< 0.001$ |
| BD | Regional | 0.08 | 0.328 |
|  | Functional epicentre | -0.15 | 0.076 |
|  | Structural epicentre | -0.33 | 0.011 |
| MDD | Regional | -0.41 | 0.005 |
| | Functional epicentre | -0.72 | $< 0.001$ |
| | Structural epicentre | -0.86 | $< 0.001$ |
| OCD | Regional | -0.13 | 0.171 |
|  | Functional epicentre | -0.38 | < 0.001 |
|  | Structural epicentre | -0.69 | < 0.001 |
| SCZ | Regional | 0.17 | 0.184 |
|  | Functional epicentre | 0.41 | 0.002 |
|  | Structural epicentre | 0.60 | < 0.001 |

ADHD, attention deficit/hyperactive disorder; ASD, autism spectrum disorder; BD, bipolar disorder; IGE, idiopathic generalized epilepsy; MDD, major depressive disorder; OCD, obsessive compulsive disorder; SCZ, schizophrenia

### Consistency of imaging-genetic analyses

Constructing PRS-HS at different P_SNP_ thresholds (*n* = 7; 0.001, 0.05, 0.1, 0.2, 0.3, 0.4, 0.5), we evaluated the robustness of the imaging-genetic effects (from ABCD, see *Figure 1* and *2*). Our findings were not affected by such methodological variations. Across the range of predictive thresholds, widespread decreases in thickness were related to PRS-HS, with strongest associations again in parietal and temporal regions (**Supplementary figure 5A**). Recapitulating the reliability of threshold-specific effects, we demonstrated high similarities among different thresholds (100.0% of correlations were significant, *p*_spin_ < 0.05). Moreover, we found comparable associations between imaging-genetic and cortical atrophy maps in left (89.2% of correlations were significant, *p*_spin_ < 0.05) and right TLE-HS (67.9% of correlations were significant, *p*_spin_ < 0.05; **Supplementary figure 5B**). Translating this approach to network models of PRS-HS, temporo-limbic and parietal epicentres identified in the main analyses were consistent across different P_SNP_ thresholds (**Supplementary figure 6A**). Moreover, the spatial distribution of these network epicentres was highly correlated with one another (100% of correlations were significant, *p*_spin_ < 0.05; **Supplementary figure 6B**).

## Discussion

Emerging literature emphasizes the importance of genotype-phenotype associations in understanding the etiological mechanisms of epilepsy. Capitalizing on recent imaging-genetic initiatives, we combined polygenic risk and whole-brain anatomy to characterize the polygenic burden of epilepsy-related HS in typical development. We found widespread decreases in cortical thickness associated with elevated PRS-HS, with the greatest effects in temporal and parietal regions. These imaging-genetic correlations were anchored to the connectivity profiles of fronto-parietal and temporo-limbic epicentres, and may play a crucial role in the network vulnerability of the brain. Structural correlates of PRS-HS mirrored case-control cortical thinning observed in patients with TLE-HS. Findings were replicable across different P_SNP_ thresholds as well as different epilepsy case-control studies. Taken together, PRS-associated structural vulnerabilities may represent an early biomarker for TLE-HS pathogenesis, offering new avenues for risk stratification and pre-emptive interventions based on their genetic profiles Structural brain organization in typical development includes a complex, and genetically determined cascade of changes from childhood to adolescence and ultimately to adulthood. Cross-sectional and longitudinal characterization of cortical gray matter tissue demonstrate global and regional thinning during this period.^38–42^ Despite being an important aspect of normal maturation, deviations from typical development have been associated with vulnerability for various neurological and psychiatric conditions, ^43–45^ including TLE-HS.^46–48^ While the exact pathogenesis of TLE-HS remains unknown, genetic studies have characterized the role of common susceptibility variants in patient cases.^13–16^ These variants account for a moderate proportion of disease phenotypic variance, and may have adverse effects on structural brain development.^15^ Core to our analytical framework is the association of individualized genetic risk profiling and mapping of structural brain phenotypes, pinpointing the morphological vulnerabilities influenced by underlying predisposition to the disease. Particularly relevant for a complex disorder that is impacted by many small-effect variants, PRS provides a personalized and compact measure of overall genetic liability.^21–23^ Linked imaging-derived phenotypes would help visualize the structural, biological impact of common variant accumulation.^28^ Examining a neurotypical population, we identified widespread cortical thinning in children with elevated PRS-HS, and conversely no relationship in the hippocampus: genetic risk is not determinant or causative of HS, but rather serves to influence the cortical alterations. Enrichment of risk variants related to focal epilepsy have been reported in patients with early onset seizures.^24,25^ Childhood-onset epilepsy has also been associated with widespread structural alterations extending beyond the seizure focus.^48,49^ Given that thickness changes in development reflect pruning and neuronal maturation,^50–52^ high genetic risk to TLE-HS may accelerate and alter synaptic elimination and/or strengthening, potentially promoting an epileptogenic network.^53^ Although no inferences can be drawn on the specific molecular mechanisms from our macroscopic imaging-genetics study, atypical structural modelling of the developing brain related to genetic risk may help predict a child’s susceptibility to epilepsy.

While imaging-genetic analyses indicate significant associations between PRS-HS and structural brain changes, the observed effect sizes are relatively low, in line with those reported in previous studies across different, genetically mediated conditions.^54–57^ It is essential to consider the context of a typically developing cohort where the genetic burden of TLE-HS is reduced. The adverse impacts of risk variants may be more subtle than those observed in a patient population with notably greater genetic vulnerability. Moreover, it is difficult to identify the mechanisms linking PRS-related morphological changes to disease onset without clear long-term follow up of epilepsy diagnosis in these healthy individuals. Patient-level data containing both genetics and imaging are necessary to address the pivot from PRS-related changes to clinically significant pathology, but have not been collected to date at a large scale. Despite these methodological challenges, using a population-based cohort, such as ABCD, provides a starting point to detect these relationships and improve our understanding of how genetic predispositions correlate with brain structural vulnerabilities.

Alterations in TLE-HS commonly implicate many brain regions organized within interconnected systems.^7,9,58–63^ Understanding these interactions and their contributions to epileptogenesis requires the integration of connectome architecture. Epicentre mapping emerges as a valuable data-driven method to pinpoint critical regions—termed epicentres—that may serve as critical anchors in the manifestation of common genetic variants.^9,33–35^ Analyzing how localized genetic vulnerabilities propagate through distributed brain regions can identify potential network pathways linking genetic risk to pathological mechanisms. In particular, marked PRS-related thinning occurs in regions strongly connected to temporo-limbic and parietal territories. Diffusion MRI is highly effective at detecting strong long-range fibre bundles and direct monosynaptic structural connections, but it does not fully capture short-range intracortical and spatially distributed polysynaptic cortical systems.^64^ By contrast, resting-state functional MRI can detect functional connectivity in the absence of direct structural connections, and thus is more informative about polysynaptic configurations.^65,66^ These temporo-limbic and parietal epicentres have been characterized by a disproportionately high number of monosynaptic and polysynaptic connections and serve as crucial areas for the integration and broadcasting of signals across different structural and functional networks. Consequently, such regions are inherently vulnerable to TLE-HS pathology.^9,62,67^ Given the convergence between functional and structural genetic epicentres, these regions also show susceptibility to the effects of accumulated genetic risk factors. Local changes related to PRS-HS may therefore impact global network organization, such that it increases vulnerability to targeted hub attacks, and potentially to seizure activity.

To bridge the transition from genetic vulnerability to clinical phenotype, we contextualized regional and network correlates of PRS with case-control atrophy and epicentres, and revealed strong spatial resemblance: thinner areas in children with elevated genetic risk tend to be thinner in patients and be highly connected to disease-related networks. Structural alterations have been consistently identified in TLE-HS, and are most marked in mesiotemporal, limbic, and sensorimotor areas.^3–7,68^ These alterations are anchored to the connectivity profiles of distinct temporo-limbic and parietal epicentres.^9^ While family-based studies have shown low heritability for these atrophy patterns.^69–71^ in healthy relatives, these predisposed regions may be too subtle and difficult to capture in endophenotype paradigms due to the complexity of epilepsy. Large sample sizes with varying genetic risk, as utilized herein, are required to characterize these imaging-genetic associations.^72^ Moreover, disease contextualization points to a common driving process between genetic risk manifestations and disease effects. As such, the polygenic burden of TLE-HS may impact biological mechanisms underlying brain structure and network architecture, and potentially influence disease vulnerability and pathogenesis. Although genetics are insufficient to cause TLE-HS alone due to its multifaceted components, they may increase susceptibility to the consequences of external factors.^20,73^ in genetically vulnerable regions and their networks.

Imaging-genetic associations also mirrored IGE-related atrophy and epicentres, to a lesser extent than in TLE-HS. Pleiotropy—whereby a genetic variant influences multiple traits–occurs in the genetics of complex traits and disorders.^74,75^ Relevant to epilepsy, certain genetic variants may contribute to the vulnerability to both generalized and focal syndromes.^15^ Despite the wide clinical spectrum of epilepsy, the shared genetic architecture may contribute to common pathological features.^76^ Supported by literature demonstrating similar patterns of cortical thinning across subtypes,^7^ our imaging-genetic model further adds to a common structural signature, such that widespread atrophy may originate from shared genetic pathways and reflect a more general epilepsy-related phenomenon. Similarly shown with disease epicentres herein, such a concept may also translate to network-level alterations. These associations may be potential biomarkers and encourage further exploration of the shared and trait-specific effects of common genetic factors in TLE-HS and the broader spectrum of epilepsy.

Limitations of imaging-genetic associations with respect to the GWAS-identified SNPs need to be highlighted. Firstly, summary statistics used for PRS calculation was based on GWAS of “focal epilepsy with documented HS”.^15^ Although it represents the most common pathological substrate for TLE-HS, hippocampal alterations occur in other epilepsy syndromes, and may be a cause, or consequence of epilepsy, or both.^10,77,78^ This phenotypic heterogeneity may impact the genetic associations identified. A more accurate delineation is crucial for detecting variants related to TLE-HS and its downstream effects, which may not be fully captured in our PRS correlations. Secondly, the same GWAS was mainly conducted in individuals of European ancestry.^15^ While our findings may be specific to European populations, they may not generalize to other under-represented groups.^79^ Replication of imaging-genetic effects, particularly using a GWAS that includes larger and more diverse cohorts—ideally with inclusion criteria that specifically define TLE-HS—could enhance the reliability and generalizability of imaging-genetic effects. This would improve the power to detect smaller effect sizes and refine the understanding of how specific genetic variants influence brain structure.

In summary, the present work highlights the potential for integrating imaging-genetic frameworks to uncover interplay between genetic predisposition, neuroanatomical changes, and epilepsy pathogenesis. Structural vulnerabilities linked to high PRS-HS in childhood resembled atrophy patterns commonly observed in patients. Collectively, these results highlight important candidates for stratification efforts that can unravel the complex etiology of epilepsy. Advancing the use of PRS as a potential biomarker for disease risk and for developing targeted interventions that prevent or limit progression of epilepsy

## Material and methods

### Participants

#### i) Adolescent Brain Cognitive Development (ABCD)

The present study used the demographic, genetic, and neuroimaging data of 3,826 unrelated neurotypical children (mean ± standard deviation [SD] age = 10.0 ± 0.6 years; 2,052 males), derived from the multi-site ABCD 2.0.1 release.^31^ Briefly, participants were recruited based on probability sampling of schools near the study sites. Parents or guardians provided written consent, while the child provided written assent. All aspects of the ABCD study were approved by the Institutional Review Board at the University of California, San Diego, United States. Overall, the large size of this cohort allows for unprecedented exploration of genetic risk for TLE-HS and its potential effects on brain organization in an *a priori* neurotypical child population.

#### ii) Human Connectome Project (HCP)

We selected 50 unrelated healthy adults. Imaging acquisition and processing are described in the **Supplementary Materials**.^80^ Such initiatives provide normative structural and functional connectivity information to employ network epicentre mapping of PRS-HS.

#### iii) Enhancing Neuro Imaging Genetics through Meta Analysis Epilepsy Consortium (ENIGMA-Epilepsy)

Imaging-genetic associations from neurotypical children were compared to MRI-based disease effects observed between 732 patients with TLE and radiological evidence of HS (mean ± SD age = 38.6 ± 10.6 years; 329 males; 391 left-sided focus) and 1,418 (mean ± SD age = 33.8 ± 10.5 years; 643 males) healthy controls (HC). Details of case-control cohorts are described in the **Supplemental Materials** and elsewhere.^28^

#### iv) Independent TLE-HS case-control datasets

To assess the replication of the aforementioned analysis, imaging-genetic associations from typically developing children were also compared to MRI-derived disease effects observed between 53 individuals with pharmaco-resistant TLE-HS and 93 age-(*t* = 1.51, *p* = 0.13) and sex-matched (χ^2^ = 0.13, *p* = 0.72) healthy controls (HC). Case-control participants were selected from (*a*) Montreal Neurological Institute and Hospital (MICs; *n*_TLE-HS/HC_ = 23/36)^29^ and (*b*) Jinling Hospital (NKG; *n*_TLE-HS/HC_ = 37/57).^30^ Sociodemographic, clinical and imaging details of patient-control cohorts are in the **Supplementary Materials**.

### Genomic data acquisition and pre-processing of ABCD

#### i) SNP Genotyping

A total of 550,000 SNPs were genotyped from saliva samples using the Illumina Human660W-Quad BeadChip. The data were prepared for imputation using “imputePrepSanger” pipeline (https://hub.docker.com/r/eauforest/imputeprepsanger/), implemented on CBRAIN^81^ and the Human660W-Quad_v1_A-b37-strand chip as reference.

#### ii) Genotyping quality control and imputation

Genotyping was quality control using PLINK 1.9.^82^ Steps were: (1) assessment of heterozygosity using the PLINK –indep-pairwise command with parameters set to 200, 50, and 0.15 to remove samples with very high or low heterozygosity; (2) removal of samples whose heterozygosity *F* coefficient was > 3 SD units from the mean; (3) removal of samples and SNPs with low call rate at 0.01 and all SNPs with minor allele frequency (MAF) < 0.01; (4) removal of individuals with mismatched sex and gender; (5) exclusion of non-European individuals by PCA with Hapmap; (6) removal of samples wih a first- or second-degree relative in the cohort (π > 0.125); (7) application of a haplotype-based test for non-random missing genotype data to remove SNPs at *p* < 1 × 10^−4^ where they had non-random associations between unobserved genotypes and missingness; and (8) application of a test for Hardy-Weinberg equilibrium (HWE) and removal of SNPs significant at *p* < 1 × 10^-6^. Imputation was performed using the Michigan Imputation Service with the Haplotype Reference Consortium (HRC) r1.1 2016 (hg19) as a reference panel.^83^

#### iii) Deriving polygenic risk scores

Individualized PRS were computed using the summary statistics from an epilepsy genome-wide association study for focal epilepsy with documented HS.^15^ While this may not necessarily equate to TLE-HS, we used this classification as a close proxy given the high prevalence and relative specificity of HS in TLE. SNPs with an INFO < 0.8 and an MAF < 0.01 were excluded, and duplicate SNPs were removed. PRSice-2 was used to calculate genetic risk scores.^84^ Given that an optimal probability threshold (P_SNP_) related to HS was previously not reported, we used multiple P_SNP_ that significantly predicted focal epilepsy: 0.001, 0.05, 0.1, 0.2, 0.3, 0.4, 0.5.^26^ All main analyses used PRS constructed at P_SNP_ < 0.1.

### Imaging acquisition and processing of ABCD

#### i) Acquisition

All participants underwent 3T MRI scanning with prospective motion correction to reduce head motion and distortions, including a 3D T1-weighted (T1w) anatomical scan based on a magnetization-prepared rapid acquisition gradient echo sequence.^31^

#### ii) Processing

T1w data were processed using FreeSurfer (version 5.3.0) to generate cortical surface and subcortical segmentations.^85,86^ Based on the Desikan-Killiany anatomical atlas,^32^ subject-specific maps of cortical thickness were sampled across 68 grey matter brain regions, and volume measures were obtained from 12 subcortical gray matter regions (bilateral amygdala, caudate, nucleus accumbens, pallidum, putamen, and thalamus) and bilateral hippocampi.

#### iii) Multisite data harmonization

Morphological data were harmonized across sites using ComBat (https://github.com/Jfortin1/ComBatHarmonization), a post-acquisition statistical batch normalization of between-site effects, while preserving age, sex and genetic risk.^87^

### Statistical analyses

#### i) Structural correlates of PRS-HS

We implemented surface-based linear models in BrainStat (version 0.4.2; https://brainstat.readthedocs.io/)^88^ with age, sex, and the first 10 genetic principal components as covariates, similar to previous imaging-genetics studies.^54–56^ These related PRS-HS to cortical thickness and subcortical volume in neurotypical children from ABCD. Multiple comparisons were corrected using the false discovery rate (FDR) procedure.^89^

#### ii) Network substrates of PRS-HS effects

We identified morphological polygenic risk epicentres by spatially correlating each brain region’s healthy functional and structural connectivity profiles from the HCP dataset to the imaging-genetic map (*i*.*e*., the unthresholded t-statistic map from *i*). This approach was repeated systematically across all cortical and subcortical regions with non-parametric spin permutation null models to control for spatial autocorrelation (5,000 repetitions),^36^ implemented in the ENIGMA toolbox (version 2.0.3; https://enigma-toolbox.readthedocs.io/).^90^ Higher the spatial similarity between a given node’s connectivity profile and whole-brain patterns of PRS-HS vulnerability supported that the node was an epicentre.

#### iii) Relation to disease-specific effects

We identified the spatial overlap between imaging-genetic alterations from ABCD and epilepsy-related alterations. The latter were obtained previously published statistical case-control atrophy and epicentre maps for left and right TLE-HS from ENIGMA-Epilepsy,^8,28^ sourced from the ENIGMA toolbox.^90^ Spin permutation-based testing (5,000 repetitions) assessed significant spatial associations between imaging-genetic and case-control effects, at the regional and network level.

We furthermore performed spatial correlations with case-control atrophy and epicentre maps for left and right TLE-HS from independent case-control datasets (MICs and NKG). Patient-specific morphology maps were *z*-scored relative to controls. We used surface-based linear models with age, sex, and site as covariates to compare between groups. Subsequent epicentre analysis identified network associations with TLE-HS atrophy patterns. Spin permutation-based testing (5,000 repetitions) evaluated significant spatial correlations between imaging-genetic and case-control effects.

To evaluate the specificity of imaging-genetic effects to TLE-HS, we repeated the same analyses with IGE^28^ and six psychiatric disorders (ADHD, ASD, BD, MDD, OCD, and SCZ), all derived from the ENIGMA Consortium.^37,90^

#### iv) Robustness analyses

To verify that results were not biased by choosing a particular threshold, we repeated the PRS analyses and associations with case-control atrophy across all predictive P_SNP_ thresholds (0.001, 0.05, 0.1, 0.2, 0.3, 0.4, 0.5).^26^ Specifically, PRS-HS was constructed at each threshold and pairwise spatial correlations between all pairs of imaging-genetic brain maps were performed. Significance testing of these correlations was assessed using spin permutation tests with 5,000 repetitions.

## Acknowledgements

AN acknowledged funding from the Fonds de la Recherche du Québec – Santé (FRQS) and the Canadian Institutes of Health Research (CIHR). SL was supported by CIHR. VK received funding from the Transforming Autism Care Consortium (TACC) and the Montreal Neurological Institute (MNI). JR received funding from the Canadian Open Neuroscience Platform (CONP) and CIHR. RRC is funded by FRQS and Healthy Brain, Healthy Lives (HBHL). ZZ received funding from the National Science Foundation of China (NSFC: 81422022; 863 project: 2014BAI04B05 and 2015AA020505) and the China Postdoctoral Science Foundation (2016M603064). LQRO and BTTY were supported by the National University of Singapore Yong Loo Lin School of Medicine (NUHSRO/2020/124/TMR/LOA), the Singapore National Medical Research Council (NMRC) Large Collaborative Grant (OFLCG19May-0035), National Medical Research Council (NMRC) Clinical Trial Grant – Investigator-Initiated Trials (CTG-IIT; CTGIIT23jan-0001), NMR Open Fund – Individual Research Grant (OF-IRG; OFIRG24jan-0030), NMRC Singapore Translational Research (STaR; STaR20nov-0003), Singapore Ministry of Health (MOH) Centre Grant (CG21APR1009), the Temasek Foundation (TF2223-IMH-01), and the National Institutes of Health (NIH; R01MH133334. BF acknowledged support from CIHR and FRQS. LC recognized funding from UNAM-DGAPA (IB201712, IG200117) and CONACYT (181508 and Programa de Laboratorios Nacionales). SSK was funded by the UK Medical Research Council (MRC) research grant (MR/S00355X/1). FC and CLY were supported by the São Paolo Research Research Foundation (FAPESP; 2013/07559-3). LB received funding from NIH and National Institute of Neurological of Neurological Disorders and Stroke (NINDS; R01NS110347). EG acknowledged support from the National Center for Advancing Translational Sciences (NCATS; UL1TR002378 and KL2TR002381). RK was funded by the Saastamoinen Foundation. TJO received support through the National Health and Medical Research Council (NHMRC) Investigator Grant (APP1176426). LV was supported by an Australian Government MRFF grants (GNT2023250, GNT1200254). SM was funded by the Ministry of Health under the Ricerca Finalizzata (NET-2013-02355313). GPW acknowledged support from the Medical Research Council (MRC; G0802012, MR/M00841X/1). ML and RG received funding from Current Research Annual Funding of the Italian Ministry of Health. KH was funded by the Heath and Care Research Wales. TR was supported by the Federal Ministry of Education and Research (epi-centre.ai, BMBF). TB received funding from the Neuro-aCSis Bonn Neuroscience Clinician Scientist Program (2024-12-07) and the BONFOR research commission of the Medical Faculty at the University of Bonn (2022-1A-21). PS was funded by the National Recovery and Resilience Plan (NRRP; PE0000006/DN.1553). JSD received support from the National Institute of Health and Care Research (NIHR) University College London Hospital (UCLH) Biomedical Research Centre. PMT was funded by R01 NS106957 and P41 EB015922. Core funding for the ENIGMA-Epilepsy Consortium was provided by the NIH Big Data to Knowledge (BD2k) program under consortium grant U54 EB020403 to PMT), CRM was supported by the NIH ( R01NS122827; R01NS124585; R01NS120976). SMS received funding from Epilepsy Society and NIH (1R01NS122827). NB and AB were funded by FRQS, CIHR, and Epilepsy Canda. ZGO received funding through grants from Michael J Fox Foundation, Canadian Consortium on Neurodegeneration in Aging Baycrest Centre for Geriatric Care, Neuro Genomics Partnership, NIH, Silverstein Foundation, Hilary and Galen Weston Foundation, and Van Berkhom Foundation. BCB acknowledged support from CIHR, SickKids Foundation, NSERC, Azrieli Center for Autism Research of the Montreal Neurological Institute (ACAR), BrainCanada, FRQS, Helmholtz International BigBrain Analytics and Learning Laboratory (HIBALL), and the Canada Research Chairs program.

## Author contributions

AN, ZGO, and BCB contributed to the conception and design of the study. AN, LL, SL, VK, SF, CW, JR, EY, RRC, ZZ, LQRO, TBTY, BF, CP, MEC, AG, LC, SSK, FC, CLY, LB, EG, NKF, RK, TJO, BS, LV, PMD, EL, AEV, SM, RK, HSZ, GPW, VJT, BAKK, ML, RG, KH, TR, OD, PS, EK, SNH, LC, MK, JSD, PMT, ENIGMA-Epilepsy, CRM, SMS, NB, AB, ZGO, and BCB contributed to the acquisition and analysis of data. AN, LL, ZGO, BCB contributed to drafting the manuscript and preparing the figures.

## Data availability

Genotyping and imaging data is available from the ABCD study upon application through NIMH Data Archive (https://nda.nih.gov/). GWAS summary statistics are available at http://www.epigad.org/gwas_ilae2018_16loci.html. The HCP dataset is available at https://db.humanconnectome.org/. Neuroimaging data from the ENIGMA-Epilepsy (meta-analysis of summary statistics) are available for download (https://github.com/MICA-MNI/ENIGMA).

## References

1. Feigin, V. L. et al. The global burden of neurological disorders: translating evidence into policy. Lancet Neurol. 19, 255–265 (2020).

2. Epilepsy, C. by H.-G. W. for the I. C. on N. of. Mesial Temporal Lobe Epilepsy with Hippocampal Sclerosis. Epilepsia 45, 695–714 (2004).

3. Lin, J. J. et al. Reduced Neocortical Thickness and Complexity Mapped in Mesial Temporal Lobe Epilepsy with Hippocampal Sclerosis. Cereb. Cortex 17, 2007–2018 (2007).

4. Keller, S. S. & Roberts, N. Voxel-based morphometry of temporal lobe epilepsy: An introduction and review of the literature. Epilepsia 49, 741–757 (2008).

5. McDonald, C. R. et al. Regional neocortical thinning in mesial temporal lobe epilepsy. Epilepsia 49, 794–803 (2008).

6. Bernhardt, B. C., Bernasconi, N., Concha, L. & Bernasconi, A. Cortical thickness analysis in temporal lobe epilepsy. Neurology 74, 1776–1784 (2010).

7. Whelan, C. D. et al. Structural brain abnormalities in the common epilepsies assessed in a worldwide ENIGMA study. Brain 141, 391–408 (2018).

8. Sisodiya, S. M. et al. The ENIGMA-Epilepsy working group: Mapping disease from large data sets. Hum. Brain Mapp. 43, 113–128 (2022).

9. Larivière, S. et al. Network-based atrophy modeling in the common epilepsies: A worldwide ENIGMA study. Sci. Adv. 6, eabc6457 (2020).

10. Lewis, D. V. et al. Hippocampal sclerosis after febrile status epilepticus: The FEBSTAT study. Ann. Neurol. 75, 178–185 (2014).

11. Hwang, S.-K. & Hirose, S. Genetics of temporal lobe epilepsy. Brain Dev. 34, 609–616 (2012).

12. EpiPM Consortium. A roadmap for precision medicine in the epilepsies. Lancet Neurol. 14, 1219–1228 (2015).

13. Kasperavičiūtė, D. et al. Epilepsy, hippocampal sclerosis and febrile seizures linked by common genetic variation around SCN1A. Brain 136, 3140–3150 (2013).

14. International League Against Epilepsy Consortium on Complex Epilepsies. Genetic determinants of common epilepsies: a meta-analysis of genome-wide association studies. Lancet Neurol. 13, 893–903 (2014).

15. Abou-Khalil, B. et al. Genome-wide mega-analysis identifies 16 loci and highlights diverse biological mechanisms in the common epilepsies. Nat. Commun. 9, 5269 (2018).

16. Stevelink, R. et al. GWAS meta-analysis of over 29,000 people with epilepsy identifies 26 risk loci and subtype-specific genetic architecture. Nat. Genet. 55, 1471–1482 (2023).

17. Kullmann, D. M. Genetics of epilepsy. J. Neurol. Neurosurg. Psychiatry 73 Suppl 2, II32–35 (2002).

18. Heyne, H. O. et al. Targeted gene sequencing in 6994 individuals with neurodevelopmental disorder with epilepsy. Genet. Med. 21, 2496–2503 (2019).

19. Mulley, J. C. et al. SCN1A mutations and epilepsy. Hum. Mutat. 25, 535–542 (2005).

20. Silvennoinen, K. et al. SCN1A overexpression, associated with a genomic region marked by a risk variant for a common epilepsy, raises seizure susceptibility. Acta Neuropathol. (Berl.) 144, 107–127 (2022).

21. Dudbridge, F. Power and Predictive Accuracy of Polygenic Risk Scores. PLOS Genet. 9, e1003348 (2013).

22. Wray, N. R. et al. Research Review: Polygenic methods and their application to psychiatric traits. J. Child Psychol. Psychiatry 55, 1068–1087 (2014).

23. Dudbridge, F. Polygenic Epidemiology. Genet. Epidemiol. 40, 268–272 (2016).

24. Gramm, M. et al. Polygenic risk heterogeneity among focal epilepsies. Epilepsia 61, e179– e185 (2020).

25. Heyne, H. O. et al. Polygenic risk scores as a marker for epilepsy risk across lifetime and after unspecified seizure events. Nat. Commun. 15, 6277 (2024).

26. Leu, C. et al. Polygenic burden in focal and generalized epilepsies. Brain 142, 3473–3481 (2019).

27. Jernigan, T. L., Brown, S. A. & Dowling, G. J. The Adolescent Brain Cognitive Development Study. J. Res. Adolesc. 28, 154–156 (2018).

28. Larivière, S. et al. Structural network alterations in focal and generalized epilepsy assessed in a worldwide ENIGMA study follow axes of epilepsy risk gene expression. Nat. Commun. 13, 4320 (2022).

29. Royer, J. et al. An Open MRI Dataset For Multiscale Neuroscience. Sci. Data 9, 569 (2022).

30. Weng, Y. et al. Macroscale and microcircuit dissociation of focal and generalized human epilepsies. Commun. Biol. 3, 1–11 (2020).

31. Casey, B. J. et al. The Adolescent Brain Cognitive Development (ABCD) study: Imaging acquisition across 21 sites. Dev. Cogn. Neurosci. 32, 43–54 (2018).

32. Desikan, R. S. et al. An automated labeling system for subdividing the human cerebral cortex on MRI scans into gyral based regions of interest. NeuroImage 31, 968–980 (2006).

33. Georgiadis, F. et al. Connectome architecture shapes large-scale cortical alterations in schizophrenia: a worldwide ENIGMA study. Mol. Psychiatry 29, 1869–1881 (2024).

34. Shafiei, G. et al. Spatial Patterning of Tissue Volume Loss in Schizophrenia Reflects Brain Network Architecture. Biol. Psychiatry 87, 727–735 (2020).

35. Zhou, J., Gennatas, E. D., Kramer, J. H., Miller, B. L. & Seeley, W. W. Predicting Regional Neurodegeneration from the Healthy Brain Functional Connectome. Neuron 73, 1216–1227 (2012).

36. Alexander-Bloch, A. F. et al. On testing for spatial correspondence between maps of human brain structure and function. NeuroImage 178, 540–551 (2018).

37. Park, B. et al. Multiscale neural gradients reflect transdiagnostic effects of major psychiatric conditions on cortical morphology. Commun. Biol. 5, 1–14 (2022).

38. Bethlehem, R. a. I. et al. Brain charts for the human lifespan. Nature 604, 525–533 (2022).

39. Giedd, J. N. et al. Brain development during childhood and adolescence: a longitudinal MRI study. Nat. Neurosci. 2, 861–863 (1999).

40. Gogtay, N. et al. Dynamic mapping of human cortical development during childhood through early adulthood. Proc. Natl. Acad. Sci. 101, 8174–8179 (2004).

41. Raznahan, A. et al. How Does Your Cortex Grow? J. Neurosci. 31, 7174–7177 (2011).

42. Shaw, P. et al. Longitudinal Mapping of Cortical Thickness and Clinical Outcome in Children and Adolescents With Attention-Deficit/Hyperactivity Disorder. Arch. Gen. Psychiatry 63, 540–549 (2006).

43. Raznahan, A. et al. A functional polymorphism of the brain derived neurotrophic factor gene and cortical anatomy in autism spectrum disorder. J. Neurodev. Disord. 1, 215–223 (2009).

44. Romero-Garcia, R., Warrier, V., Bullmore, E. T., Baron-Cohen, S. & Bethlehem, R. A. I. Synaptic and transcriptionally downregulated genes are associated with cortical thickness differences in autism. Mol. Psychiatry 24, 1053–1064 (2019).

45. Valk, S. L., Di Martino, A., Milham, M. P. & Bernhardt, B. C. Multicenter mapping of structural network alterations in autism. Hum. Brain Mapp. 36, 2364–2373 (2015).

46. Beghi, E. The Epidemiology of Epilepsy. Neuroepidemiology 54, 185–191 (2019).

47. Bozzi, Y., Casarosa, S. & Caleo, M. Epilepsy as a Neurodevelopmental Disorder. Front. Psychiatry 3, (2012).

48. Hermann, B. et al. The Neurodevelopmental Impact of Childhood-onset Temporal Lobe Epilepsy on Brain Structure and Function. Epilepsia 43, 1062–1071 (2002).

49. Boutzoukas, E. M. et al. Cortical thickness in childhood left focal epilepsy: Thinning beyond the seizure focus. Epilepsy Behav. 102, 106825 (2020).

50. Huttenlocher, P. R. & Dabholkar, A. S. Regional differences in synaptogenesis in human cerebral cortex. J. Comp. Neurol. 387, 167–178 (1997).

51. Paus, T. Growth of white matter in the adolescent brain: Myelin or axon? Brain Cogn. 72, 26–35 (2010).

52. Rakic, P.Bourgeois, J.-P. & Goldman-Rakic, P. S. Synaptic development of the cerebral cortex: implications for learning, memory, and mental illness. in Progress in Brain Research (eds. Van Pelt, J., Corner, M. A., Uylings, H. B. M. & Lopes Da Silva, F.H.) vol. 102 227– 243 (Elsevier, 1994).

53. Goldberg, E. M. & Coulter, D. A. Mechanisms of epileptogenesis: a convergence on neural circuit dysfunction. Nat. Rev. Neurosci. 14, 337–349 (2013).

54. de Mol, C. L. et al. Polygenic Multiple Sclerosis Risk and Population-Based Childhood Brain Imaging. Ann. Neurol. 87, 774–787 (2020).

55. Khundrakpam, B. et al. Neural correlates of polygenic risk score for autism spectrum disorders in general population. Brain Commun. 2, fcaa092 (2020).

56. Kirschner, M. et al. Schizophrenia Polygenic Risk During Typical Development Reflects Multiscale Cortical Organization. Biol. Psychiatry Glob. Open Sci. 3, 1083–1093 (2023).

57. Stauffer, E.-M. et al. Grey and white matter microstructure is associated with polygenic risk for schizophrenia. Mol. Psychiatry 26, 7709–7718 (2021).

58. Bernhardt, B. C., Chen, Z., He, Y., Evans, A. C. & Bernasconi, N. Graph-Theoretical Analysis Reveals Disrupted Small-World Organization of Cortical Thickness Correlation Networks in Temporal Lobe Epilepsy. Cereb. Cortex 21, 2147–2157 (2011).

59. Bernhardt, B. C. et al. The spectrum of structural and functional imaging abnormalities in temporal lobe epilepsy. Ann. Neurol. 80, 142–153 (2016).

60. Crossley, N. A. et al. The hubs of the human connectome are generally implicated in the anatomy of brain disorders. Brain 137, 2382–2395 (2014).

61. Hatton, S. N. et al. White matter abnormalities across different epilepsy syndromes in adults: an ENIGMA-Epilepsy study. Brain 143, 2454–2473 (2020).

62. Royer, J. et al. Epilepsy and brain network hubs. Epilepsia 63, 537–550 (2022).

63. Yasuda, C. L. et al. Aberrant topological patterns of brain structural network in temporal lobe epilepsy. Epilepsia 56, 1992–2002 (2015).

64. Maier-Hein, K. H. et al. The challenge of mapping the human connectome based on diffusion tractography. Nat. Commun. 8, 1349 (2017).

65. Benkarim, O. et al. A Riemannian approach to predicting brain function from the structural connectome. NeuroImage 257, 119299 (2022).

66. Honey, C. J. et al. Predicting human resting-state functional connectivity from structural connectivity. Proc. Natl. Acad. Sci. 106, 2035–2040 (2009).

67. Gleichgerrcht, E. et al. Temporal Lobe Epilepsy Surgical Outcomes Can Be Inferred Based on Structural Connectome Hubs: A Machine Learning Study. Ann. Neurol. 88, 970–983 (2020).

68. Bonilha, L. et al. Extrahippocampal gray matter loss and hippocampal deafferentation in patients with temporal lobe epilepsy. Epilepsia 51, 519–528 (2010).

69. Alhusaini, S. et al. Temporal Cortex Morphology in Mesial Temporal Lobe Epilepsy Patients and Their Asymptomatic Siblings. Cereb. Cortex 26, 1234–1241 (2016).

70. Alhusaini, S. et al. Normal cerebral cortical thickness in first-degree relatives of temporal lobe epilepsy patients. Neurology 92, e351–e358 (2019).

71. Yaakub, S. N. et al. Abnormal temporal lobe morphology in asymptomatic relatives of patients with hippocampal sclerosis: A replication study. Epilepsia 60, e1–e5 (2019).

72. Westlye, L. T., Alnæs, D., van der Meer, D., Kaufmann, T. & Andreassen, O. A. Population-Based Mapping of Polygenic Risk for Schizophrenia on the Human Brain: New Opportunities to Capture the Dimensional Aspects of Severe Mental Disorders. Biol. Psychiatry 86, 499–501 (2019).

73. Perucca, P. & Scheffer, I. E. Genetic Contributions to Acquired Epilepsies. Epilepsy Curr. 21, 5–13 (2021).

74. Gratten, J. & Visscher, P. M. Genetic pleiotropy in complex traits and diseases: implications for genomic medicine. Genome Med. 8, 78 (2016).

75. Sivakumaran, S. et al. Abundant Pleiotropy in Human Complex Diseases and Traits. Am. J. Hum. Genet. 89, 607–618 (2011).

76. Leu, C. et al. Pleiotropy of polygenic factors associated with focal and generalized epilepsy in the general population. PLOS ONE 15, e0232292 (2020).

77. Kim, H., Mansi, T. & Bernasconi, N. Disentangling Hippocampal Shape Anomalies in Epilepsy. Front. Neurol. 4, (2013).

78. Thom, M. Review: Hippocampal sclerosis in epilepsy: a neuropathology review. Neuropathol. Appl. Neurobiol. 40, 520–543 (2014).

79. Peterson, R. E. et al. Genome-wide Association Studies in Ancestrally Diverse Populations: Opportunities, Methods, Pitfalls, and Recommendations. Cell 179, 589–603 (2019).

80. Van Essen, D. C. et al. The WU-Minn Human Connectome Project: An overview. NeuroImage 80, 62–79 (2013).

81. Sherif, T. et al. CBRAIN: a web-based, distributed computing platform for collaborative neuroimaging research. Front. Neuroinformatics 8, (2014).

82. Chang, C. C. et al. Second-generation PLINK: rising to the challenge of larger and richer datasets. GigaScience 4, s13742-015-0047–8 (2015).

83. McCarthy, S. et al. A reference panel of 64,976 haplotypes for genotype imputation. Nat. Genet. 48, 1279–1283 (2016).

84. Choi, S. W. & O’Reilly, P. F. PRSice-2: Polygenic Risk Score software for biobank-scale data. GigaScience 8, giz082 (2019).

85. Dale, A. M., Fischl, B. & Sereno, M. I. Cortical Surface-Based Analysis: I. Segmentation and Surface Reconstruction. NeuroImage 9, 179–194 (1999).

86. Fischl, B. FreeSurfer. NeuroImage 62, 774–781 (2012).

87. Fortin, J.-P. et al. Harmonization of cortical thickness measurements across scanners and sites. NeuroImage 167, 104–120 (2018).

88. Larivière, S. et al. BrainStat: A toolbox for brain-wide statistics and multimodal feature associations. NeuroImage 266, 119807 (2023).

89. Benjamini, Y. & Hochberg, Y. Controlling the False Discovery Rate: A Practical and Powerful Approach to Multiple Testing. J. R. Stat. Soc. Ser. B Methodol. 57, 289–300 (1995).

90. Larivière, S. et al. The ENIGMA Toolbox: multiscale neural contextualization of multisite neuroimaging datasets. Nat. Methods 18, 698–700 (2021).

