## Supplementary Materials for "Associations between epilepsy-related polygenic risk and brain morphology in childhood"

**Supplementary material and methods**

**Human connectome project dataset**

*i) MRI acquisition*. Functional and diffusion data of 50 unrelated healthy adults was obtained from the Human Connectome Project (HCP) 3T database. MRI parameters details can be found elsewhere^1^. Given its high-quality imaging data and its common use in the generation of normative connectomes, the HCP dataset provides a reliable reference for the network epicentre mapping.

*ii) MRI processing*. Multimodal processing utilized *micapipe* (version 0.2.3; <https://micapipe.readthedocs.io/>).^2^ T1-weighted (T1w) data were deobliqued, reoriented to standard neuroscience orientation (LPI: left to right, posterior to anterior, and inferior to superior), corrected for intensity nonuniformity, and skull-stripped. FreeSurfer (version 6.0.0) was used to generate models of the inner and outer cortical interfaces.^3,4^ Subcortical structures were segmented using FSL FIRST.^5^ Native diffusion weighted images (DWI) were denoised, underwent b0 intensity normalization and were corrected for susceptibility distortion, head motion and eddy currents. Resting-state functional MRI processing included slice-timing and head motion correction. Nuisance and global signals were removed. Preprocessed timeseries were parcellated according to the Desikan-Killiany anatomical atlas.^6^

*iii) Connectome generation*. Normative functional connectivity matrices were generated by computing pairwise correlations between the timeseries of all 68 cortical regions and between all subcortical and cortical regions; negative connections were set to zero. Subject-specific connectivity matrices were then *z*-transformed and aggregated across participants to construct a group-average functional connectome.

We generated normative structural connectivity matrices from preprocessed DWI using MRtrix3.^7,8^ Anatomically constrained tractography was performed using different tissue types (cortical and subcortical grey matter, white matter, cerebrospinal fluid) derived from subject-specific T1w images.^9^ Multishell and multi-tissue response functions were estimated, and constrained spherical deconvolution and intensity normalization was performed. Tractograms were generated with 40 million streamlines (maximum tract length = 250; fractional anisotropy cut-off = 0.06). We applied spherical deconvolution informed filtering of tractograms (SIFT2) to reconstruct whole brain streamlines weighted by the cross-sectional multipliers. Reconstructed streamlines were projected onto the 68 cortical, 12 subcortical, and hippocampal areas. Individual-specific structural connectivity matrices were log transformed to reduce connectivity strength variance and further averaged across participants.

**Epilepsy case-control datasets: ENIGMA-Epilepsy**

*i) Participants*. Imaging-genetic analyses were correlated to disease effects in 732 individuals diagnosed with TLE and radiological evidence of HS (mean ± SD age = 38.6 ± 10.6 years; 329 males; 391 left-sided focus) and 1418 site-matched healthy control (mean ± SD age = 33.8 ± 10.5 years; 643 males).^10,11^ Patients were diagnosed according to the seizure and syndrome classifications of the ILAE, namely the combination of electroclinical features and MRI findings typically associated with underlying HS. Individuals with a progressive or neurodegenerative disease, malformations of cortical development, tumors, or prior neurosurgery were excluded. Furthermore, controls had no history of mental disorders. Local institutional review boards and ethics committees approved each included cohort study, and written informed consent was provided according to local requirements. Further details on specific case-control cohorts are referenced elsewhere.^10^

*ii) MRI acquisition*. Participants underwent T1w brain MRI at each centre. Scanner and acquisition protocols detailed elsewhere.^10,12^

*iii) MRI processing*. Using the standard ENIGMA workflow, images were processed at each site. Models of cortical surfaces were generated with FreeSurfer (version 5.3.0)^3^ and cortical thickness was measured based on the Desikan-Killiany anatomical atlas.^6^ Missing data were imputed with the mean value for that given region; participants with missing data in at least half of the cortical or subcortical brain measures were excluded.

*iv) Multisite data harmonization*. Morphological data were harmonized across sites using ComBat (<https://github.com/Jfortin1/ComBatHarmonization>), a post-acquisition statistical batch normalization to harmonize between-site effects while preserving effects of age, sex and disease status.^13^

**Epilepsy case-control datasets: MICs and NKG**

*i) Participants*. We replicated our correlation analyses with 53 individuals with pharmaco-resistant TLE-HS and 93 age- (*t* = 1.51, *p* = 0.13) and sex-matched (*χ*² = 0.13, *p* = 0.72) healthy controls (HC). Case-control participants were selected from two independent sites: (a) Montreal Neurological Institute and Hospital (MICs; *n*_TLE/HC_ = 23/36)^14^ and (b) Jinling Hospital (NKG; *n*_TLE/HC_ = 37/57).^15^ Detailed sociodemographic and clinical information of patient-control cohorts are noted in **Supplementary table 1**. Patients were diagnosed according to the classification of the International League Against Epilepsy based on a comprehensive evaluation including clinical history, seizure semiology, video-electroencephalography recordings, neuroimaging, and/or neuropsychological assessment. We excluded patients who had encephalitis, a history of traumatic brain injury, or bilateral TLE diagnosis. At the time of data analysis, all patients had standard radiologically suspected hippocampal sclerosis (HS) from preoperative examination, with 46 having undergone resective surgery (40 had undergone cortico-amygdalo-hippocampectomy and 6 had undergone selective amygdalo-hippocampectomy). At post-surgical follow-up (mean ± standard deviation [SD] = 2.42 ± 2.18 years), 34 patients have been seizure-free (Engel I), 5 have shown significant reductions in seizure frequency (Engel II), 4 have shown prolonged seizure intervals (Engel III), 2 have shown no worthwhile improvements (Engel IV), and 1 was lost for follow-ups. Based on established histopathological criteria, all available specimens showed HS or gliosis. Data collection was approved by the ethics committees of each site, and all participants provided written informed consent in accordance with the Declaration of Helsinki.

*ii) MRI acquisition*. Every participant underwent a research-dedicated multimodal MRI scan before surgery: (a) Data from Montreal Neurological Institute and Hospital (MICs) were collected on a 3T Siemens Magnetom Prisma-Fit scanner equipped with a 64-channel head coil, and included a 3D-MPRAGE (voxel size=0.8×0.8×0.8mm3, matrix size=320×320, repetition time [TR]=2300ms, echo time [TE]=3.14ms, flip angle [FA]=9°, inversion time [TI]=900ms, 224 slices); (b) Data from Jinling Hospital (NKG) were collected on a 3T Siemens Trio scanner with a 32-channel head coil, and included a 3D-MPRAGE (voxel size=0.5×0.5×1mm3, TR=2300ms, TE=2.98ms, FOV=256×256mm2, FA=9°).

*iii) MRI processing*. All Native T1w structural images were deobliqued, reoriented, intensity nonuniformity corrected, skull-stripped, and submitted to FreeSurfer (version 6.0.0)  to extract surface models of the cortical mental.^3,4^ Subject-specific cortical thickness was measured across all Desikan-Killiany parcels.^6^

*iv) Multisite data harmonization*. Morphological data were harmonized across sites using ComBat (<https://github.com/Jfortin1/ComBatHarmonization>) while preserving effects of age, sex and disease status.^13^

**References**

1.	Glasser, M. F. *et al.* The minimal preprocessing pipelines for the Human Connectome Project. *NeuroImage* **80**, 105–124 (2013).

2.	Cruces, R. R. *et al.* Micapipe: A pipeline for multimodal neuroimaging and connectome analysis. *NeuroImage* **263**, 119612 (2022).

3.	Dale, A. M., Fischl, B. & Sereno, M. I. Cortical Surface-Based Analysis: I. Segmentation and Surface Reconstruction. *NeuroImage* **9**, 179–194 (1999).

4.	Fischl, B. FreeSurfer. *NeuroImage* **62**, 774–781 (2012).

5.	Jenkinson, M., Beckmann, C. F., Behrens, T. E. J., Woolrich, M. W. & Smith, S. M. FSL. *NeuroImage* **62**, 782–790 (2012).

6.	Desikan, R. S. *et al.* An automated labeling system for subdividing the human cerebral cortex on MRI scans into gyral based regions of interest. *NeuroImage* **31**, 968–980 (2006).

7.	Tournier, J.-D. *et al.* *MRtrix3*: A fast, flexible and open software framework for medical image processing and visualisation. *NeuroImage* **202**, 116137 (2019).

8.	Tournier, J.-D., Calamante, F. & Connelly, A. MRtrix: Diffusion tractography in crossing fiber regions. *Int. J. Imaging Syst. Technol.* **22**, 53–66 (2012).

9.	Smith, R. E., Tournier, J.-D., Calamante, F. & Connelly, A. Anatomically-constrained tractography: Improved diffusion MRI streamlines tractography through effective use of anatomical information. *NeuroImage* **62**, 1924–1938 (2012).

10.	Larivière, S. *et al.* Structural network alterations in focal and generalized epilepsy assessed in a worldwide ENIGMA study follow axes of epilepsy risk gene expression. *Nat. Commun.* **13**, 4320 (2022).

11.	Sisodiya, S. M. *et al.* The ENIGMA-Epilepsy working group: Mapping disease from large data sets. *Hum. Brain Mapp.* **43**, 113–128 (2022).

12.	Whelan, C. D. *et al.* Structural brain abnormalities in the common epilepsies assessed in a worldwide ENIGMA study. *Brain* **141**, 391–408 (2018).

13.	Fortin, J.-P. *et al.* Harmonization of cortical thickness measurements across scanners and sites. *NeuroImage* **167**, 104–120 (2018).

14.	Royer, J. *et al.* An Open MRI Dataset For Multiscale Neuroscience. *Sci. Data* **9**, 569 (2022).

15.	Weng, Y. *et al.* Macroscale and microcircuit dissociation of focal and generalized human epilepsies. *Commun. Biol.* **3**, 1–11 (2020).

**Supplementary tables and figures**

**Supplementary table 1 | Demographic and clinical breakdown of epilepsy case-control cohort**

| **Demographics** | **MICs** | | **NKG** | |
| --- | --- | --- | --- | --- |
|  | **HC** | **TLE** | **HC** | **TLE** |
| Number | 36 | 16 | 57 | 37 |
| Age, y | 35.5(9.2) | 44.4(11.3) | 25.6(5.9) | 26.7(7.7) |
| Sex, M/F | 16/20 | 7/9 | 25/32 | 18/19 |
| Focus side, L/R | – | 9/7 | – | 16/21 |
| Onset age, y | – | 24.1(17.9) | – | 14.9(7.9) |
| Duration, y | – | 20.3(14.6) | – | 11.8(7.1) |
| Engel I/II/III/IV | – | 8/0/0/0 | – | 26/5/4/2 |
| HS^1^ | – | 9/9 | – | 37/37 |

HC, healthy control; TLE, temporal lobe epilepsy; M, male; F, female; L, left; R, right; y, years; Age, age at seizure onset, and epilepsy duration are presented as mean (±SD) years. ^1^Histologically confirmed HS in available post-surgical specimens.

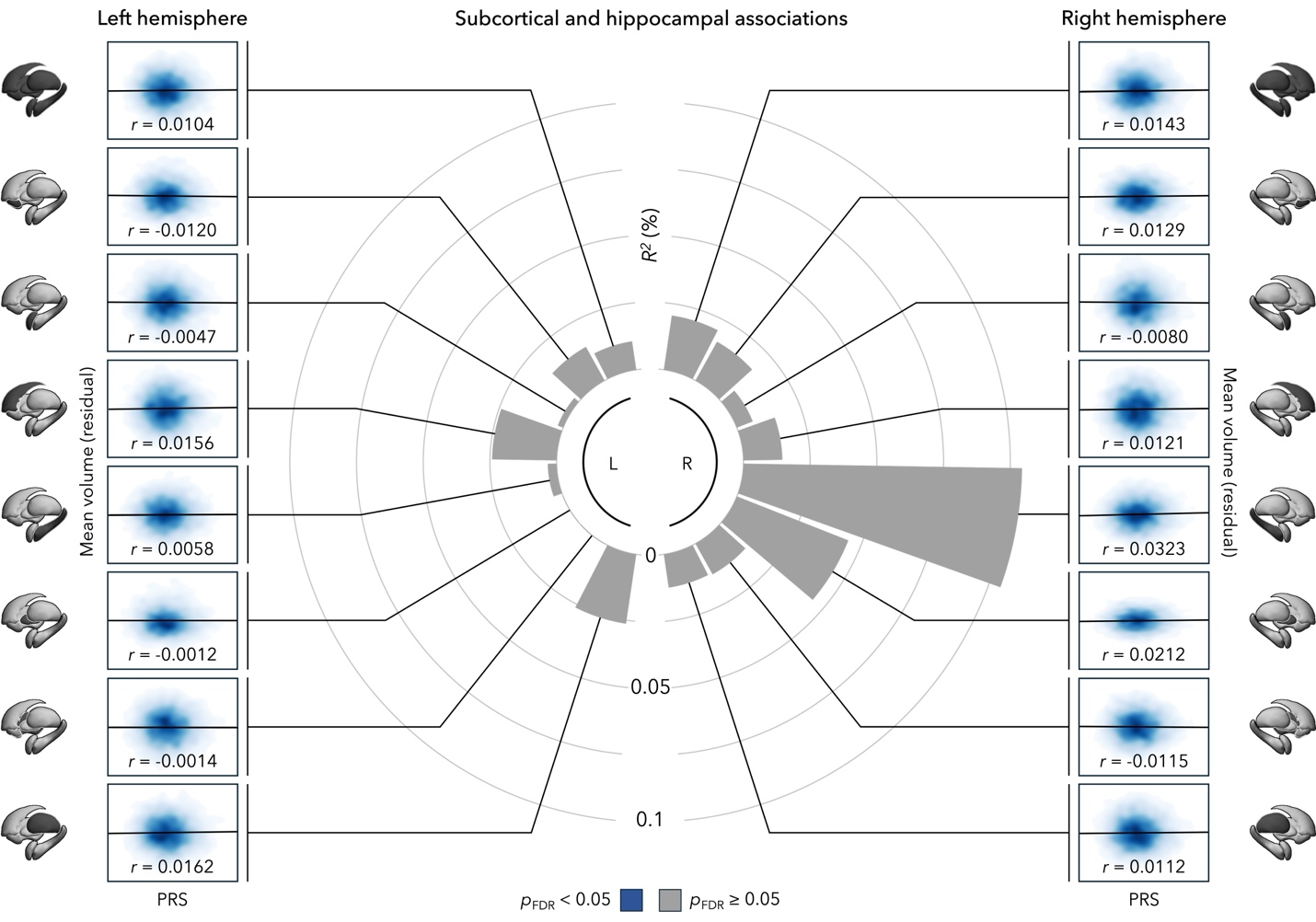

**Supplementary figure 1 | PRS-HS associations with subcortical and hippocampal volume (ABCD).** Distribution of genetic risk effects on morphology across the different regions (in order from top to bottom: all, accumbens, amygdala, caudate, hippocampus, pallidum, putamen, thalamus). L, left; PRS-HS; polygenic risk score for epilepsy-related hippocampal sclerosis; R, right.

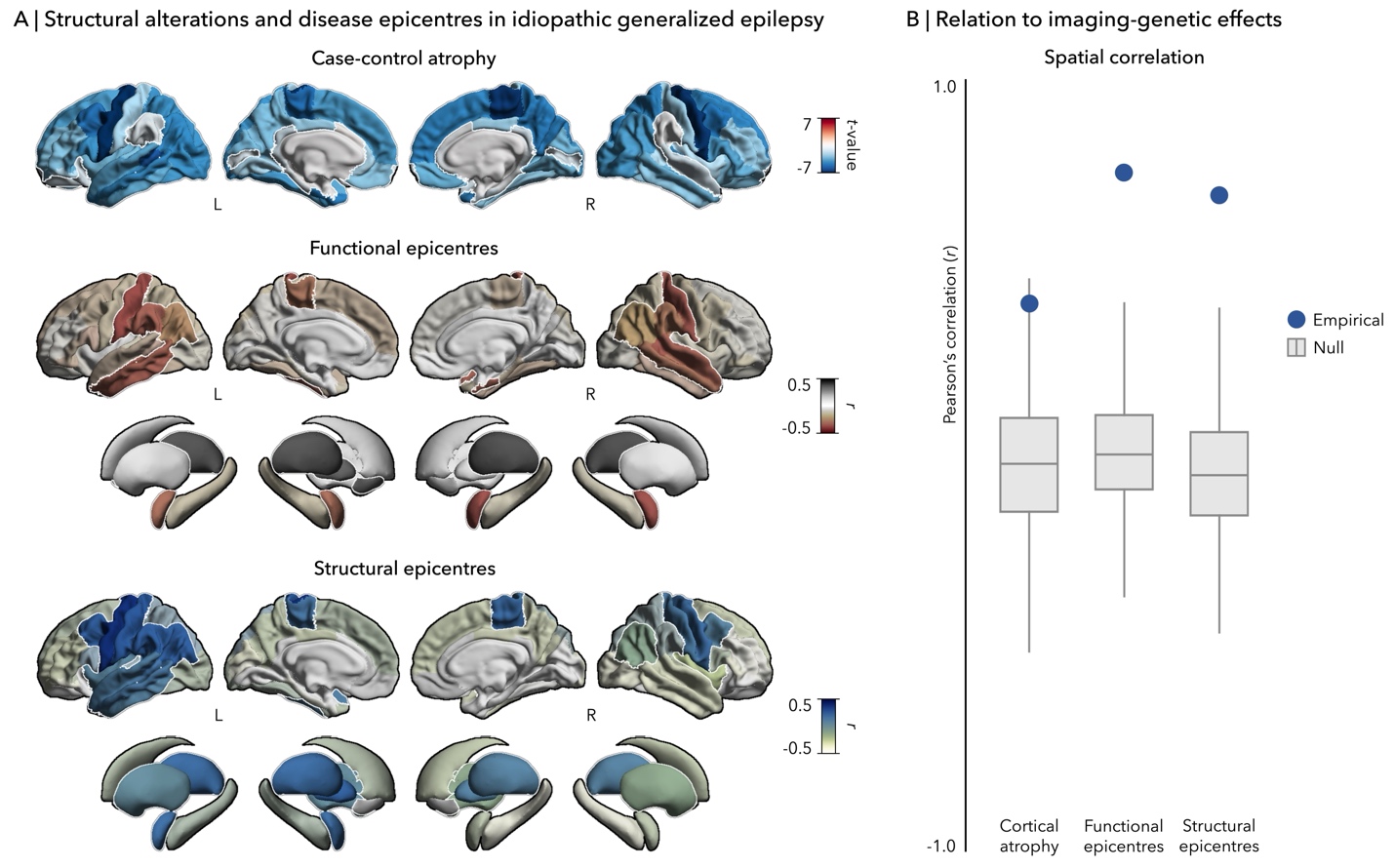

**Supplementary figure 2 | Comparison between imaging-genetic (ABCD) and IGE-related disease effects (ENIGMA). (A)** Cortical atrophy and disease epicentres in IGE. Outline in white represents *p*_FDR/spin_ < 0.05. IGE, idiopathic generalized epilepsy; L, left; R, right. **(B)** Spatial correlations between genetic and disease maps are compared against permutation-based null correlations. Points represents the empirical correlation (red and blue ponts depict *p*_spin_ < 0.05). In the boxplots, the ends of boxes represent the first (25%) and third (75%) quartiles, the centre line (median) represents the second quartile of the null distribution (*n* = 5,000 permutations), the whiskers represent the non-outlier endpoints of the distribution.

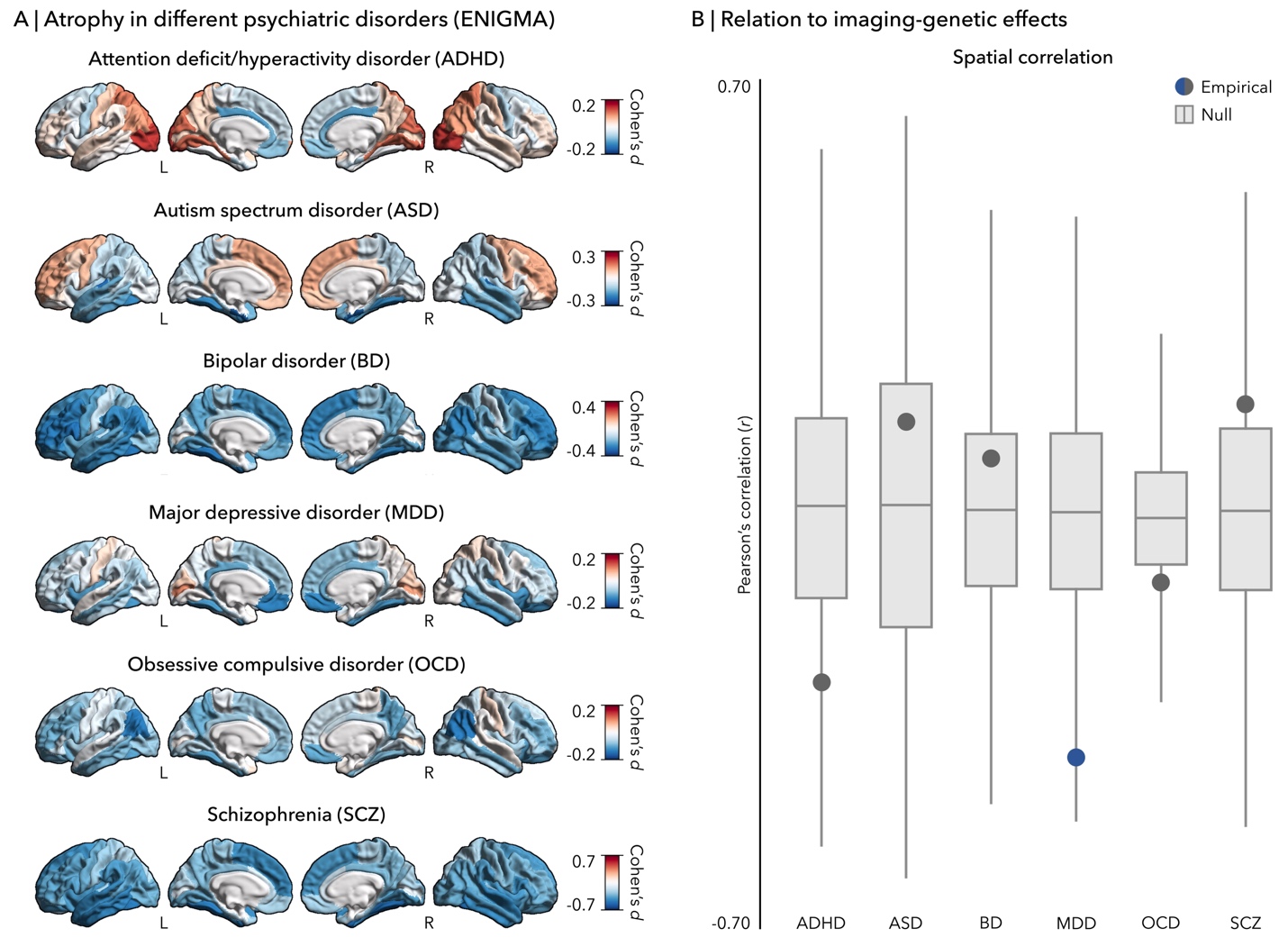

**Supplementary figure 3 | Comparison between PRS-HS effects (ABCD) and psychiatric disorder atrophy (ENIGMA). (A)** Case-control differences in attention deficit/hyperactivity disorder (ADHD), autism spectrum disorder (ASD), bipolar disorder (BD), major depressive disorder (MDD), obsessive compulsive disorder (OCD), and schizophrenia (SCZ).  Blue and red colours point to atrophy and hypertrophy in patients relative to healthy controls, respectively. Outline in white represents *p*_FDR_ < 0.05. L, left; R, right. (B) Spatial correlations between structural alterations in psychiatric disorders and imaging-genetic maps are compared against permutation-based null correlations. Points represent the empirical correlation (blue points signify *p*_spin_ < 0.05). In the boxplots, the ends of boxes represent the first (25%) and third (75%) quartiles, the centre line (median) represents the second quartile of the null distribution (*n* = 5,000 permutations), the whiskers represent the non-outlier endpoints of the distribution.

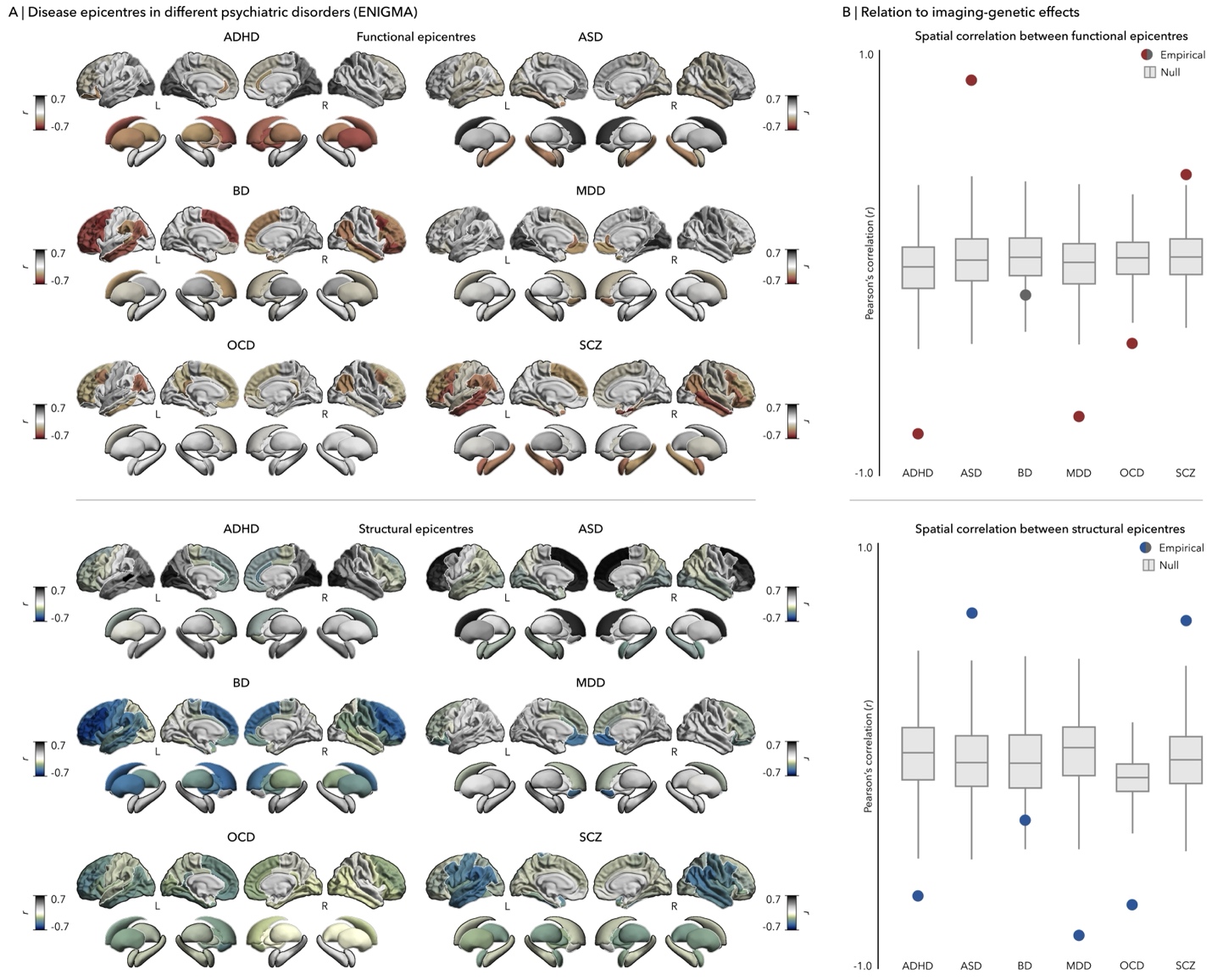

**Supplementary figure 4 | Comparison between imaging-genetic (ABCD) and psychiatric disease epicentres (ENIGMA). (A)** Functional and structural disease epicentres in ADHD, ASD, BD, MDD, OCD, SCZ.  Red and blue colours represent negative associations, while grey depicts positive correlations. Outline in white represents *p*_spin_ < 0.05. ADHD, attention deficit/hyperactivity disorder; ASD, autism spectrum disorder; BD, bipolar disorder; L, left; MDD, major depressive disorder; OCD, obsessive compulsive disorder; SCZ, schizophrenia; R, right; TLE-HS, temporal lobe epilepsy with hippocampal sclerosis. (B) Spatial correlations between genetic and disease epicentre maps are compared against permutation-based null correlations. Points represent the empirical correlation (red and blue points depict *p*_spin_ < 0.05). In the boxplots, the ends of boxes represent the first (25%) and third (75%) quartiles, the centre line (median) represents the second quartile of the null distribution (*n* = 5,000 permutations), the whiskers represent the non-outlier endpoints of the distribution.

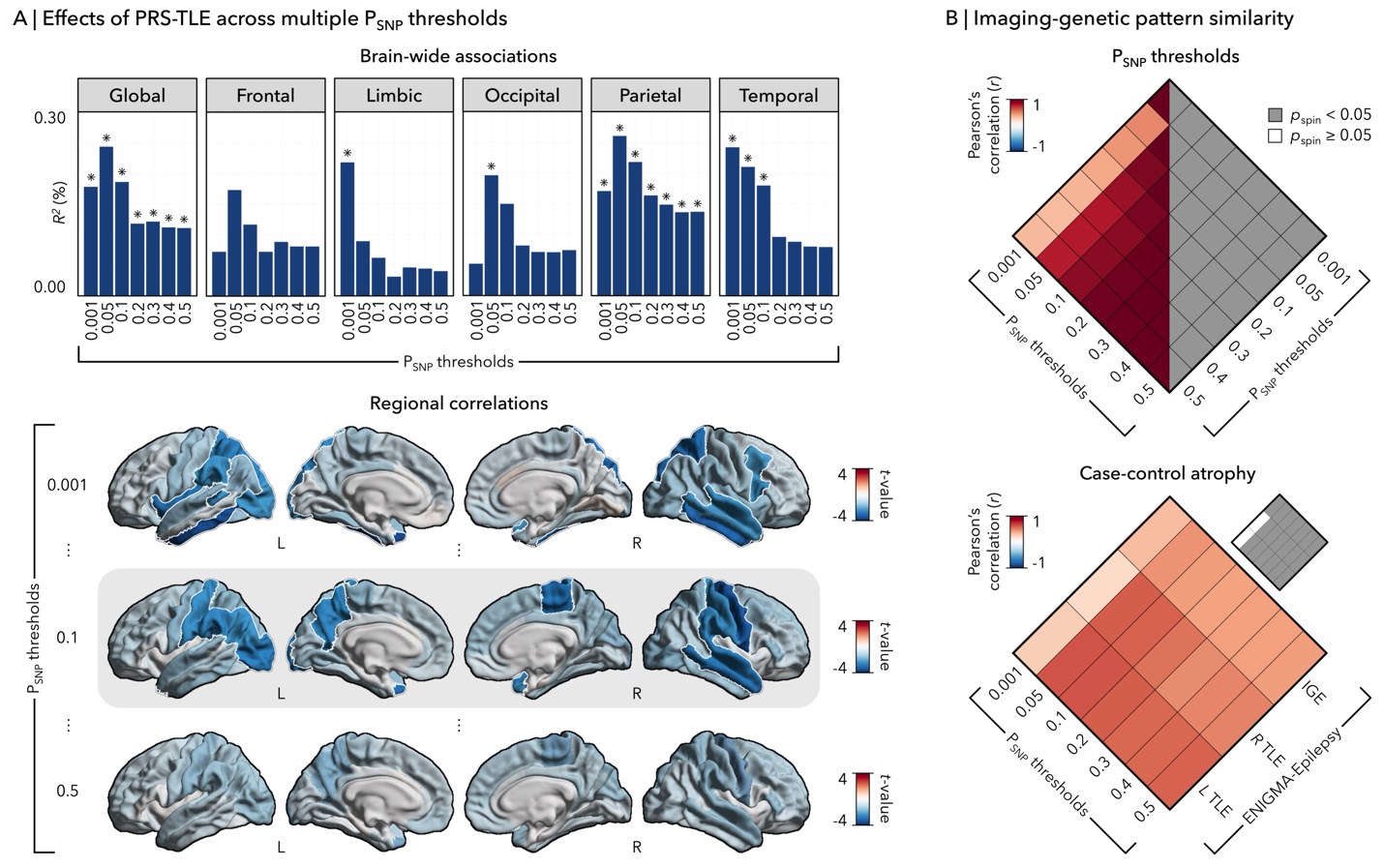

**Supplementary figure 5 | Region-level consistency of PRS-HS effects at different P_SNP_ thresholds (ABCD). (A)** PRS-related changes in cortical thickness at the global (top) and regional (bottom) level. White outline indicates significant correlations (*p*_FDR_ < 0.05). L, left; PRS-HS; polygenic risk score for hippocampal sclerosis; R, right; SNP, single nucleotide polymorphism. **(B)** Associations between imaging-genetic correlations across the different P_SNP_ thresholds (top) and with ENIGMA-Epilepsy case-control atrophy (bottom). Statistical significance was assessed using two-tailed, non-parametric tests.

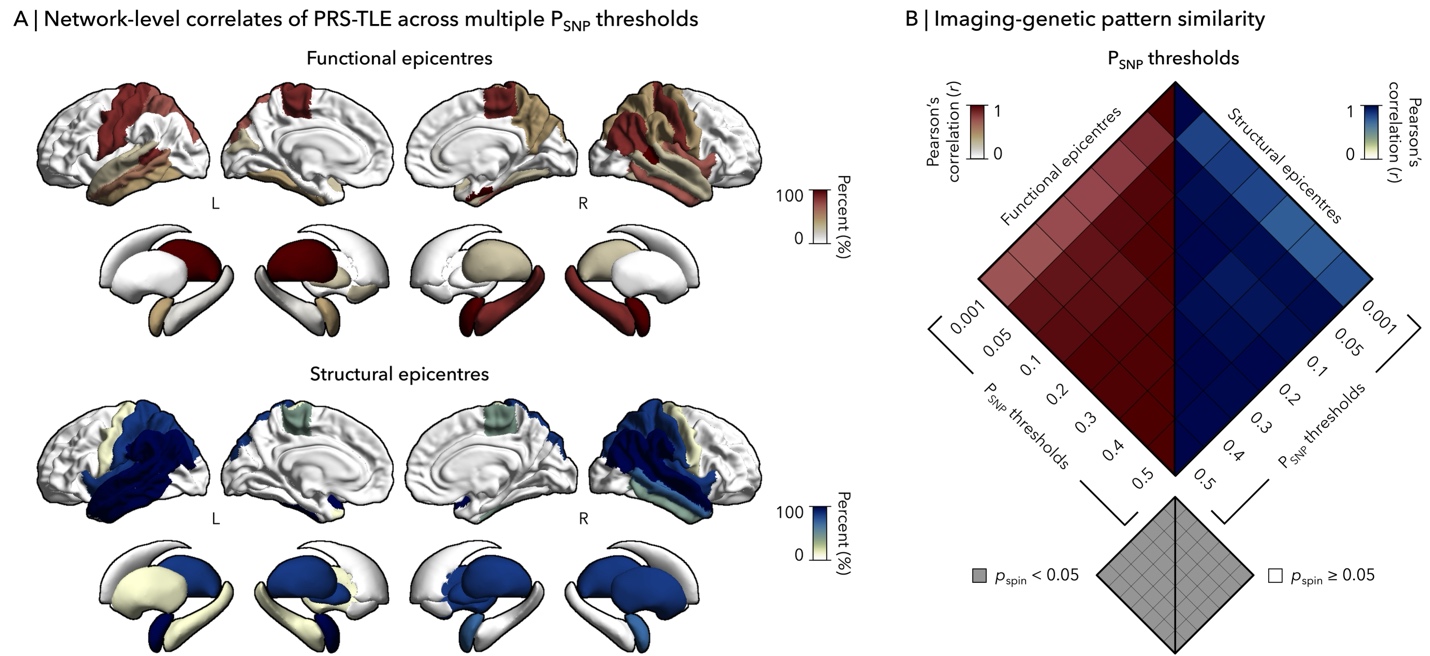

**Supplementary figure 6 | Network-level consistency of PRS-HS effects at different PSNP thresholds (ABCD). (A)** Percentage of thresholds showing significant functional (top) and structural (bottom) epicentres across different thresholds. L, left; PRS-HS; polygenic risk score for epilepsy-related hippocampal sclerosis; R, right; SNP, single nucleotide polymorphism. **(B)** Spatial correlation between all pairs of thresholds. Statistical significance was assessed using two-tailed, non-parametric tests.
